# Outgrowth of Mouse Sensory Neurons is not Mediated by Regeneration Associated Genes in Early Postnatal Stages

**DOI:** 10.64898/2026.10.06.757081

**Authors:** Beatriu Molina-Esteve, Jose Crugeiras, María Rodríguez-Brañas, Ana Sanchez-Maldonado, David Ovelleiro, Natalia Lago, Esther Udina

**Affiliations:** Medical Physiology Unit, Faculty of Medicine, Cell Biology, Physiology and Immunology and Institute of Neurosciences, Universitat Autònoma de Barcelona Bellaterra, Spain; Centro de Investigación Biomédica en Red sobre Enfermedades Neurodegenerativas (CIBERNED), Bellaterra, Spain; Peripheral Nervous System, Vall d’Hebron Institut de Recerca (VHIR), Vall d’Hebron Hospital Universitari, Vall d’Hebron Barcelona Hospital Campus, Barcelona, Spain

**Keywords:** Neonatal nerve injury, Nerve regeneration, Regeneration Associated Genes, Transcriptomic analysis, Conditioning lesion

## Abstract

Peripheral neurons exhibit robust regeneration after nerve injury in adulthood; a process linked to the activation of regeneration-associated genes (RAGs). However, the mechanisms underlying nerve regeneration during early postnatal development remain poorly understood. In a previous work, we observed that motoneurons axotomized at early postnatal stages regenerate without overexpressing RAGs, contrasting with the expected RAG activation observed in postnatal day 30 (P30) mice. To determine whether this phenomenon extends to sensory neurones, we investigated two paradigmatic dorsal root ganglia populations: proprioceptive (PV) and nociceptive (TrpV1) neurones, which exhibit distinct regenerative capacities in adult mice.

Following sciatic nerve crush injury at P4, P10, and P30, both neuronal subtypes demonstrated robust axonal regeneration across all postnatal stages, with nociceptors exhibiting significantly superior regeneration compared to proprioceptors. Remarkably, no substantial neuronal death was observed at any developmental timepoint. However, RNA sequencing revealed that P4 and P10 sensory neurones failed to upregulate RAGs following injury, whereas P30 neurones exhibited the expected pro-regenerative transcriptional programme. Pathway enrichment analysis confirmed that classical regeneration-associated signalling pathways were activated exclusively in young adults, whilst developmental and inflammatory pathways predominated in early postnatal stages.

Using the conditioning lesion paradigm in primary dorsal root ganglia cultures, we demonstrated that only P30 neurones exhibited enhanced neurite outgrowth following conditioning lesion, whilst P4 and P10 neurones showed no conditioning effect or reduced outgrowth, respectively. These findings indicate that early postnatal sensory neurones possess intrinsic growth capacity independent of RAG activation, suggesting fundamentally distinct regenerative mechanisms across developmental stages.

**Graphical abstract:** 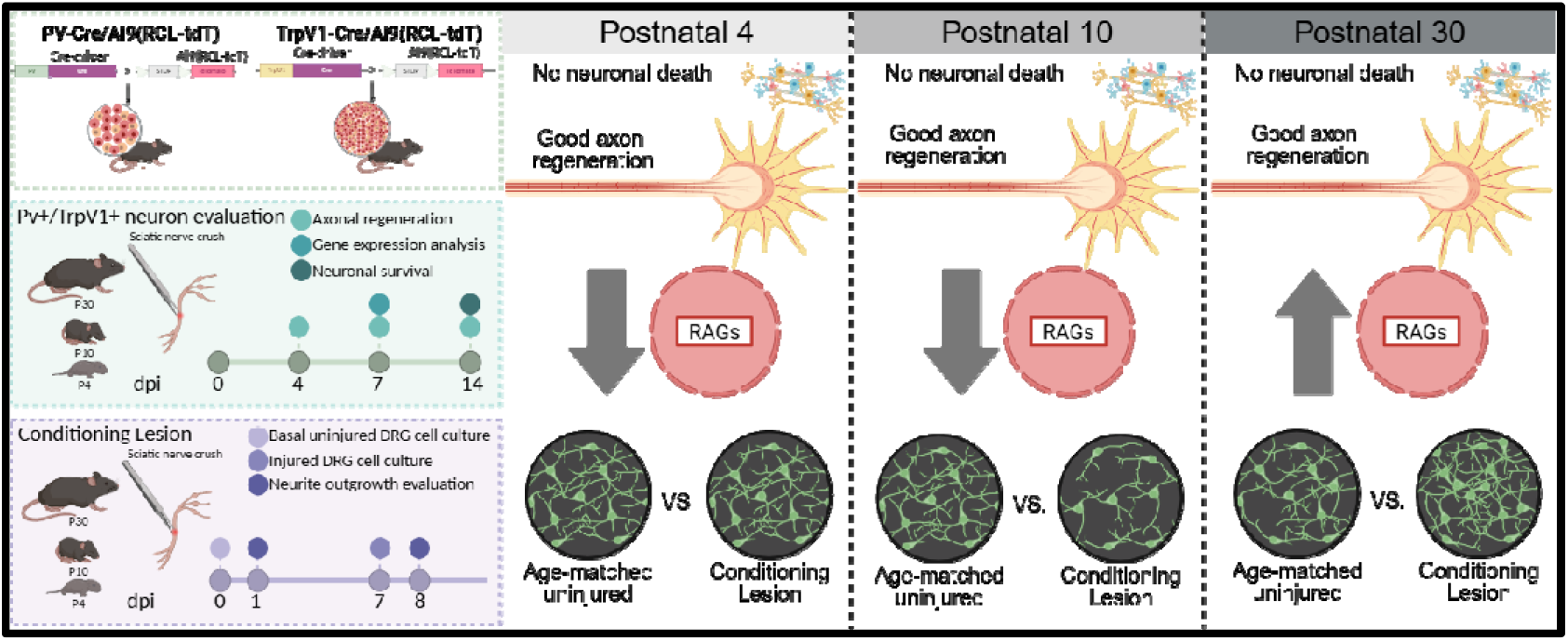

## Introduction

It is well known that peripheral neurons can regenerate after nerve injuries. This regenerative ability has been linked to a successful Wallerian degeneration in the distal nerve stump to create a permissive environment for growth, and to the capacity of injured neurons to switch from a neurotransmitter state to a pro-regenerative state, that includes the overexpression of regeneration associated genes (RAGs), key elements for a successful regeneration. However, this phenomenon does not occur in a uniform way across all neuronal subpopulations (Navarro, Verdú, and Butí 1994; Janko 1976; Brushart et al., 2020; Bolívar et al., 2024) nor during all developmental stages (Kemp et al., 2015a; Kemp et al., 2015b).

During the postnatal stage, the peripheral nervous system undergoes profound transformations that influence neuronal plasticity and the response to injury. In contrast to adulthood, young neurons display a greater susceptibility to neuronal death following axotomy (Lowrie and Vrbová 1984; Lowrie et al., 1987; Pollin et al., 1991) yet they also possess a molecular environment that can promote axonal regrowth and functional recovery, provided they survive the injury (Donnerer 2003; Christie and Zochodne 2013; Renthal et al., 2020). Despite that classical works focus mainly on the massive death of motoneurons after postnatal nerve injuries, sensory neurons can also be affected (Kemp et al., 2015a).

In a previous work (Molina-Esteve et al., 2025) we analysed the regenerative capacity of motoneurons after injuries at different postnatal stages, and we observed that despite a good regenerative response, motoneurons axotomized at postnatal day 4 (P4) or postnatal day 10 (P10) did not overexpress RAGS, whereas postnatal day 30 (P30) mice showed the expected activation of these pro-regenerative genes. To further elucidate the growth programs activated by postnatal neurons after axotomy, we have explored the regenerative capability of two paradigmatic sensory subtypes (proprioceptors and nociceptors) at different postnatal stages. Within the dorsal root ganglia (DRG), proprioceptive neurons (PV+) constitute a large-diameter neuronal population characterized by myelinated axons that innervate muscle spindles and Golgi tendon organs. They play a crucial role in proprioception and the regulation of motor coordination (Madden et al., 2020; Wu et al., 2021). Nociceptive sensory neurons (TrpV1^+^) are distinguishable by their small soma diameters and unmyelinated (C fibres) or lightly myelinated axons (Madden et al., 2020) and are responsible for the chemical, thermic and mechanic nociception (Dubin and Patapoutian 2010).

These two populations show different regenerative capability in adult mice (Bolívar et al., 2024). The genetic program that these populations or motoneurons activated after axotomy is also different, but share common pathways, that include the overexpression of RAGs (Bolívar et al., 2024). Therefore, in this work we want to investigate the regenerative response of proprioceptive and nociceptive sensory neurons, whether postnatal sensory neurons neither express RAGs when injured at early postnatal stages and whether a prior lesion to the sciatic nerve is conditioning the neuron to promote regeneration.

As expected, both proprioceptive and nociceptive neurons showed good regenerative potential when injured at different postnatal stages, being nociceptors significatively better than proprioceptors. However, an RNAseq of the whole DRG showed that P4 and P10 sensory neurons did not overexpress RAGs after injury, as we had previously observed in injured motoneurons of the same age. To further elucidate whether these postnatal neurons do not overexpress RAGs because they still retain a growth state or they activate a different pro-regenerative program, we performed primary sensory cultures and used the conditioning lesion paradigm (Smith and Skene 1997). Since neurons need to switch to a pro-regenerative program to be able to extend neurites, a previous axotomy would positively condition the neurons. This conditioning lesion effect would not be present in postnatal neurons if their growth ability is due to their basal state and not to the activation of a strong pro-regenerative program as the one observed in adults.

## Material and Methods

### Mice

P4 to P30-day-old mice were used for all studies. Males and females were used in approximately equal numbers for all groups in all experiments. The mice were housed in a controlled environment (12-h light-dark cycle, 22 ± 2°C) in open cages with water and food *ad libitum*. P4 and P10 mice were housed with their mothers. The strains used in this study were acquired from Jackson Laboratory (Bar Harbor, ME, USA) and maintained in our animal facility. C57BL/6J mice were used as wild type (WT) animals. Mice expressing the fluorescent protein TdTomato in the proprioceptive neurons Pv-Cre/Ai9 and nociceptive neurons TrpV1-cre/Ai9 were generated by breeding homozygous Ai9 (RCL-tdT) mice (JAX stock #007909) (Madisen et al., 2010) with B6 PV-Cre (Parvalbumin, JAX stock #017320) or TRPV1^cre^ (transient receptor potential cation channel, subfamily V, member 1, JAX stock #017769).

### Experimental design

For the evaluation of axonal regeneration and neuronal death, a sciatic nerve crush injury was applied to PV-Cre/Ai9(RCL-tdT) and TrpV1-Cre/Ai9(RCL-tdT) mice at P4, P10 and P30, followed by histological analyses (Figure 1a). To characterize the transcriptomics of the sensory neurons after a crush injury at the same postnatal stages, RNA from L3-L5 DRGs WT mice was isolated and RNAseq was performed (Figure 1b). To assess if a conditioning lesion promotes neurite outgrowth in vitro, a sciatic nerve crush was applied to WT mice at the same postnatal stages, and neurons from L3-L5 DRGs were cultivated for 24 hours (Figure 1c).

**Figure 1.**
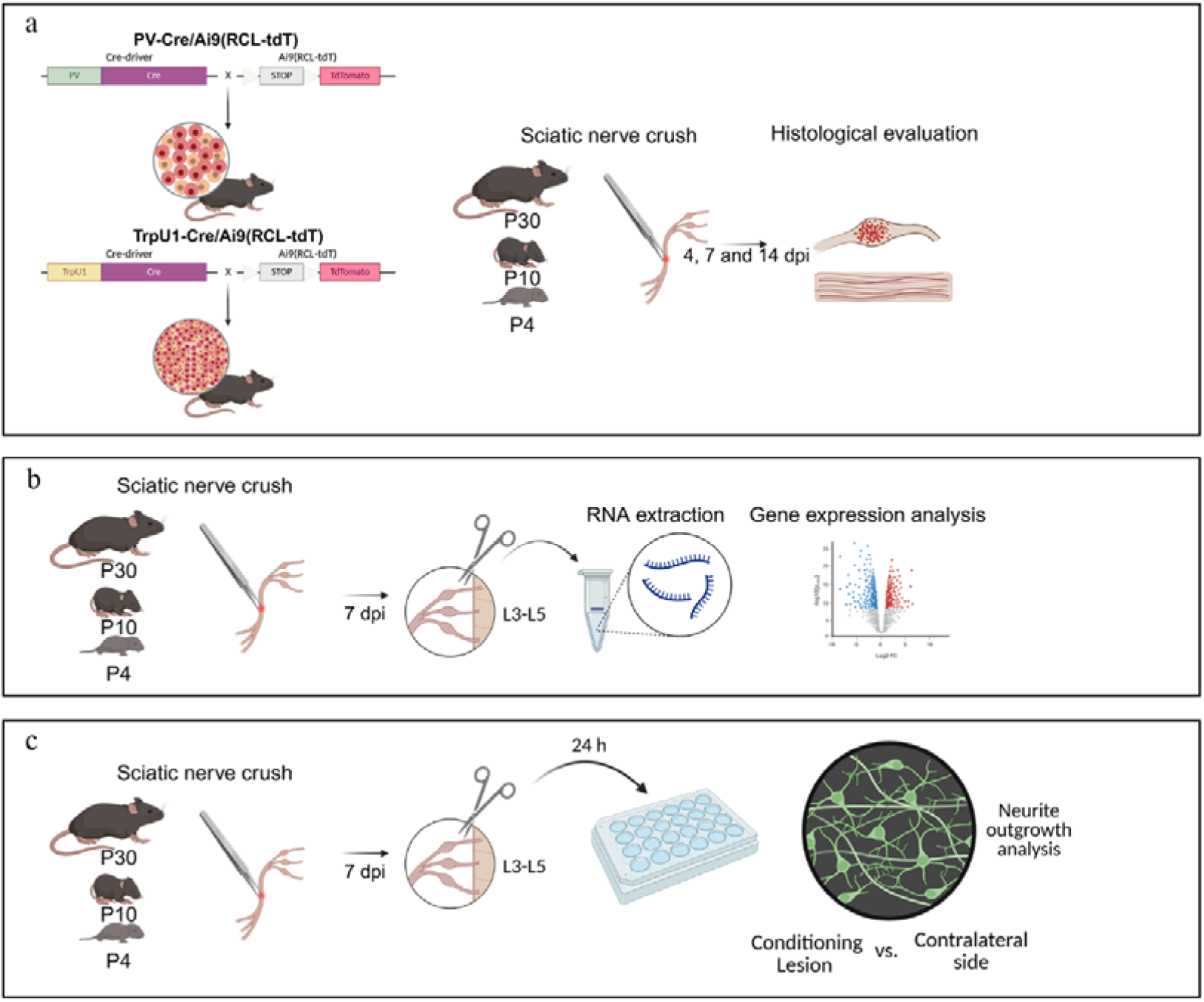
Experimental design (a) Schematic representation of the transgenic mice and experimental design used for the evaluation of proprioceptive and nociceptive sensory neurons survival and axonal regeneration. (b) Schematic representation of the experimental design used for sensory neurons transcriptomics analysis. (c) Schematic representation of the experimental design used for evaluation of neurite outgrowth after a conditioning lesion. Intact animals were also used to evaluate basal growth.

### Surgeries

All experimental procedures were approved by the Animal Experimentation Ethical Committee of the Universitat Autònoma de Barcelona and followed the European Communities Council Directive 2010/63/EU and Spanish National law (RD 53/2013). All surgical procedures were carried out aseptically under an operating microscope. Ketamine (90 mg/kg or 18mg/kg) and xylazine (10 mg/kg or 2 mg/kg) was administered intraperitoneally to anaesthetize P30 and P10 mice respectively. In P4 animals, short-term hypothermia was used to anaesthetize the pups, taking advantage of the short period needed to perform the surgery (Flecknell et al., 2015). A mid-thigh incision was made, and the right sciatic nerve was exposed and crushed. Standard N°. Five Jeweller forceps were used for 30 seconds, resulting in axonotmesis or Sunderland grade 2 injury (Bridge et al., 1994). The incision was sutured with a 6–0 silk suture and disinfected with povidone iodine. All mice were kept in a warm environment until recovery. During the follow-up, assessment of general health and mobility was performed daily for the first week. All the animals were sacrificed at the time of study termination under deep anaesthesia with Dolethal (0.03 ml/30 g) administered intraperitoneally. Death was confirmed by decapitation.

### Axon regeneration and neuronal death

To evaluate the number of proprioceptive and nociceptive axons per nerve and the number of neuron cell bodies 4, 7 and 14 days after sciatic nerve crush injury, the ipsilateral and contralateral sciatic nerves and the ipsilateral and contralateral L3-L5 DRGs were harvested and stored in PBS with 30% sucrose at 4°C, being post-fixed before storage. The sciatic nerves were placed on a glass slide and mounted with Fluoromount-G medium (Southern Biotech). Pressure was applied to the coverslips to flatten the nerves, and images 5 and 10 mm distal from the lesion point were taken via a confocal microscope (Leica SP5, 20x, z-step size of 1 μm). Axons were counted in stacks of 5 μm every 10 μm to avoid double counting and compared to the contralateral side. The L3-L5 DRGs were placed on a glass slide and mounted with Fluoromount-G medium (Southern Biotech). Images every 1 μm were taken via a confocal microscope (Leica SP5, 20x, z-step size of 1 μm). Proprioceptive and nociceptive neurons were counted in stacks of 1 μm every 20 μm to avoid double counting and compared to its contralateral side to assess neuronal death. All neuron counts were corrected with the Abercrombie formula (Abercrombie and Johnson 1946) *(N=nt/(t+d))* where *N* is the corrected number of counted neurons, *n* is the number of cells that have been counted, *t* is the thickness of the section, and *d* is the average diameter of the cells. The diameter of neurons was measured in 100 randomly chosen neurons (per group), using Image Tool software and the mean value was calculated. To calculate the percentage of neuronal death or neuronal survival, the total number of neurons on the ipsilateral side was compared with the total number of neurons on the contralateral side.

### Transcriptomic analyses

Seven days after the sciatic nerve crush, all mice were euthanized with intraperitoneal Dolethal (0.03 ml/30 g) and the L3-L5 DRGs were dissected and placed on cold’s Gey’s solution enriched with glucose (6 mg/mL) until used. DRG’s were homogenized in 1 mL of buffer. Both female and male were used in this experiment, at least four animals were used for age and group. After centrifugation of the homogenate, 50 μL of the supernatant was used for RNA isolation using the RNAeasy Micro Kit (QIAGEN, #74004) and quantified wit Quant-it RiboGreen RNA assay kit (Thermo Fisher, #R11490).

#### Library preparation and RNA sequencing

BGI Genomics Co., Ltd. (Shenzhen, China) performed the RNA quality control and RNA sequencing. Library preparation and RNA sequencing for this report were carried out as follows: The library type was “DNBSEQ Eukaryotic Strand-specific mRNA library,” and all samples were sequenced on the DNBSEQ platform (DNBSEQ Technology) with a read length of 2x150 bp (PE150), ensuring a minimum of 20 million clean reads per sample, always reaching Q20 and Q30 quality scores above 95% and 89%, respectively. Library quality control included mRNA enrichment from total RNA using oligo dT, mRNA fragmentation, and double-stranded cDNA synthesis using random primers, with dUTP incorporation in the second strand to ensure strand specificity. Next, end-repair and 3’ adenylation, adapter ligation, PCR, library quality control, circularization, and amplification in DNA nanoballs were performed before sequencing. FASTQ file quality was assessed using SOAPnuke software (developed by BGI), and the results confirm that the data meets quality and filtering standards, using Phred+33 encoding. Reads were quantified using the coding DNA reference database (Ensembl Mouse database, genome assembly: GRCm39.cdna.all, release 113) via Salmon (v1.10.3). A total of 27 samples were analysed. The "DESeq2" library (v 1.48.0) was utilised to perform the differential expression analysis pipeline. Only genes with more than 10 counts in the smallest group were included in the differential analysis. Principal Component Analysis (PCA) was performed to visualise the overall effect of experimental covariates and batch effects. Uniform Manifold Approximation and Projection (UMAP) was implemented with n_neighbors parameter set to 15. Hierarchical clustering was performed using the R "pheatmap" function (v 1.0.12), focusing on the 80 most variable genes determined using the MatrixGenerics::rowVars function. Gene set enrichment was performed using the R library "STRINGdb" (v 2.20) via the STRING platform. Only genes with P-values<0.05 were included in the analysis. Multiple databases were utilised, including Gene Ontology (Biological Process, Molecular Function, Cellular Component), PubMed reference publications, KEGG pathways and Reactome pathways among others. Pathway analysis was conducted using Bioconductor libraries "KEGGREST" (v 1.48) and "pathview" (v 1.48). The KEGG database was accessed via RESTful web services through the R package.

### In vitro cultures of DRG sensory neurons

Cell cultures of the L3-L5 DRGs were performed in intact P4, P10 and P30 mice or mice at these postnatal stages one week after receiving a sciatic nerve crush (conditioning lesion). Mice were euthanized administering intraperitoneally an overdose of Dolethal (0.03 ml/30 g), and the L3-L5 DRGs were dissected and placed in colds Gey’s solution (Sigma, Cat #G9779) enriched with glucose (6 mg/mL) (Sigma, Cat #G7021-1KG) until used. Bilateral DRG were used in intact animals. In animals previously injured, homolateral lumbar DRG were used to evaluate the conditioning lesion effect, being the contralateral used as a control. DRGs were enzymatically dissociated in Hank’s Balanced Salt Solution (Sigma, Cat #14170-088) supplemented with 10% trypsin (2.5 mg/mL; Sigma, Cat #T-4674), 10% collagenase A (12.5 collagen digestion units/ml; Sigma, Cat #C2674) and 10% DNAse (1 mg/mL; Roche, Cat #11284932001) for 45 minutes at 37°C, followed by mechanical dissociation. After the enzymes were blocked with 10% high Foetal Bovine Serum (hiFBS) in Dulbecco’s Modified Eagle Medium (DMEM) (Sigma, Cat #41966052) and centrifuged at 900 rpm for 7 min, the cells were centrifuged again in 15% Bovine Serum Albumin (BSA) (Sigma, Cat #A6003) in Neurobasal A (Gibco, Cat #10088022). The supernatant was withdrawn, and 1 mL of culture medium was added. The medium consisted of Neurobasal A supplemented with 2% B27 (Gibco, Cat #17504044), 0.033 M glucose, 1 mM glutamine (Gibco, Cat #35050-038), 1× penicillin/streptomycin (Sigma, Cat #P0781), and 20 mM HEPES (Sigma, Cat #H0887). To culture the cells, 24-well plates were used. The plates were previously coated with 100 μg/mL poly-D-lysine (PDL; Sigma, Cat #P6407) for 2 h at 37°C, washed, dried and further coated with 1 μL/mL laminin for a minimum of 2 hours (Sigma, Cat #L-2020). Then, neurons were plated at a density of 3 × 10^3^ neurons/mL and incubated for 24h.

Cell cultures were fixed with 4% paraformaldehyde (PFA) in phosphate buffer saline (PBS) buffer for 20 minutes, then washed three times with PBS and incubated for 30 minutes in blocking solution (5% normal donkey serum (NDS) in PBS-Triton 0,3% (PBST)). After removal of the blocking solution, cultures were incubated for 1 hour at room temperature in a humidified chamber with the primary antibody mouse anti-β-III-tubulin (1:500, Biolegend, #MMS435P) diluted in blocking solution. Following three washes in PBS, cultures were incubated for 2 hours at room temperature with the appropriate secondary antibody (Alexa Fluor 488 Donkey x Mouse, Invitrogen, A21202) diluted in blocking solution and protected from light. Three additional PBS washes were performed. Finally, coverslips were mounted onto glass slides using Fluoromount G mounting medium (Southern Biotech, #0100-20) and allowed to dry before imaging.

#### Neurite outgrowth quantification

The samples were examined using a fluorescence microscope (Nikon Eclipse Ni, Japan) equipped with a DS-RI2 digital camera and operated via the dedicated NIS Elements software. Image acquisition was performed for each well using a 10x objective, with a resolution of 1600×1200 pixels. For the morphometric quantification of neurite growth, the obtained images were processed and analysed with Fiji/ImageJ software. The analysis included the delineation and measurement of the longest neurite per neuron, using a manual tool within the software.

### Data analysis

All statistical analyses were performed using GraphPad Prism 9 (version 9.0.1). The normality of data distributions was assessed with the Shapiro-Wilk test (p>0.05 was considered normally distributed). Data are presented as mean ± standard error of the mean (SEM). Statistical significance was set at p<0.05. For comparisons of multiple groups and variables, two-way ANOVA was used, where appropriate, post hoc Sidak’s correction was applied for pairwise multiple comparisons. For paired measurements, paired t-tests were applied and one-way ANOVA with Tukey’s post hoc test was employed for single-factor group comparisons. When the data did not meet criteria for normal distribution, non-parametric analysis was performed using the Kruskal-Wallis’s test followed by Dunn’s test. For transcriptomic data, differential gene expression was analysed using DESeq2 after filtering for minimum read counts. Principal Component Analysis (PCA), Uniform Manifold Approximation and Projection (UMAP), and hierarchical clustering were used for exploratory and batch effect analyses. Gene set enrichment and pathway analyses were conducted using STRINGdb and Bioconductor libraries.

## Results

### Response of proprioceptive and nociceptive neuros to nerve injury at different postnatal stages

#### Evaluation of neuronal somas

Neuronal survival was assessed 14 dpi by quantifying the total number of neuronal cell bodies in the ipsilateral (L3–L5) DRGs. Pv-Cre/Ai9(RCL-tdT) mice were used to identify proprioceptive neurons (Figure 2a), and TrpV1-Cre/Ai9(RCL-tdT) mice to identify nociceptive neurons (Figure 2c), across P4, P10, and P30 animals. Due to high variability between animals and the differences in the size of ganglia between ages, all comparisons were performed within the same animal (using the contralateral lumbar DRG).

**Figure 2.**
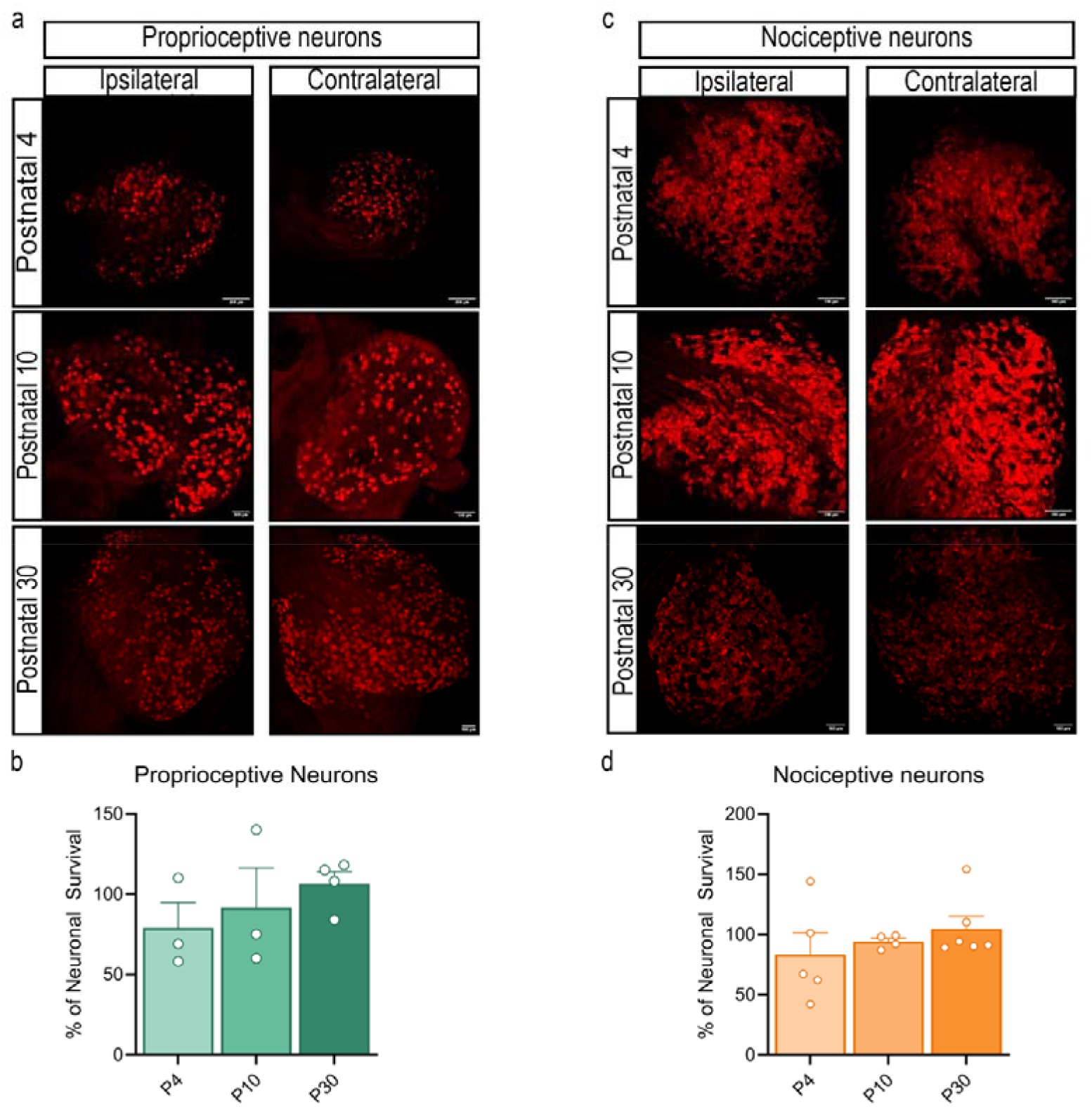
Survival of proprioceptive and nociceptive neurons after a nerve injury at different postnatal stages. (a) Representative images of the ipsilateral and contralateral L3-L5 DRGs of P4 mice at 14 dpi, showing the PV^+^ neurons in red. Scale bar= 200 µm. (b) Percentage of surviving PV^+^ neurons at 14 dpi in the different postnatal stages. Statistical analysis was performed via one-way ANOVA followed by Tukey’s correction test. (c) Representative images of the ipsilateral and contralateral L3-L5 DRGs of P4 mice at 14 dpi, showing the TrpV1^+^ neurons in red. Scale bar= 200 µm. (d) Percentage of surviving TrpV1^+^ neurons at 14 dpi in the different postnatal stages. Statistical analysis was performed via one-way ANOVA followed by Tukey’s correction test.

The large diameter and low number of proprioceptive neurones facilitate the counting and about 300 neurons in total were counted (Supplementary Figure 1). We estimated a survival rate of ∼80% at P4 and >90% at P10 and P30 (Figure 2b) reduction that was not significant at any age (p>0.05). The smaller diameter and large number of nociceptive neurons difficult the counting and lead to high variability between ages, but percentage of survival was similar to the one reported in proprioceptors, of about ∼80% at P4 and reached ∼100% at P10 and P30 (Figure 2d). No significant neuronal loss was detected at any age (p>0.05).

#### Evaluation of axonal regeneration

When analysing the number of proprioceptive and nociceptive axons in intact nerves at different postnatal stages (Figures 3, 4), we found a significantly higher number of PV^+^ axons in P10 and P30 mice compared with P4 mice (p<0.05; Figure 4a). Since PV^+^ axons are highly myelinated, incomplete myelination at early postnatal stage could difficult the visualization of these axons in P4 animals in the confocal microscope. For TrpV1^+^ axons, P10 mice showed a greater number of axons than P4 mice (p<0.05), although this difference was not observed when compared with P30 mice (p>0.05; Figure 4e). Again, the lower amount of myelin in P4 nerves could reduce the number of axons detected by the confocal microscopy at this age compared to more mature animals.

**Figure 3.**
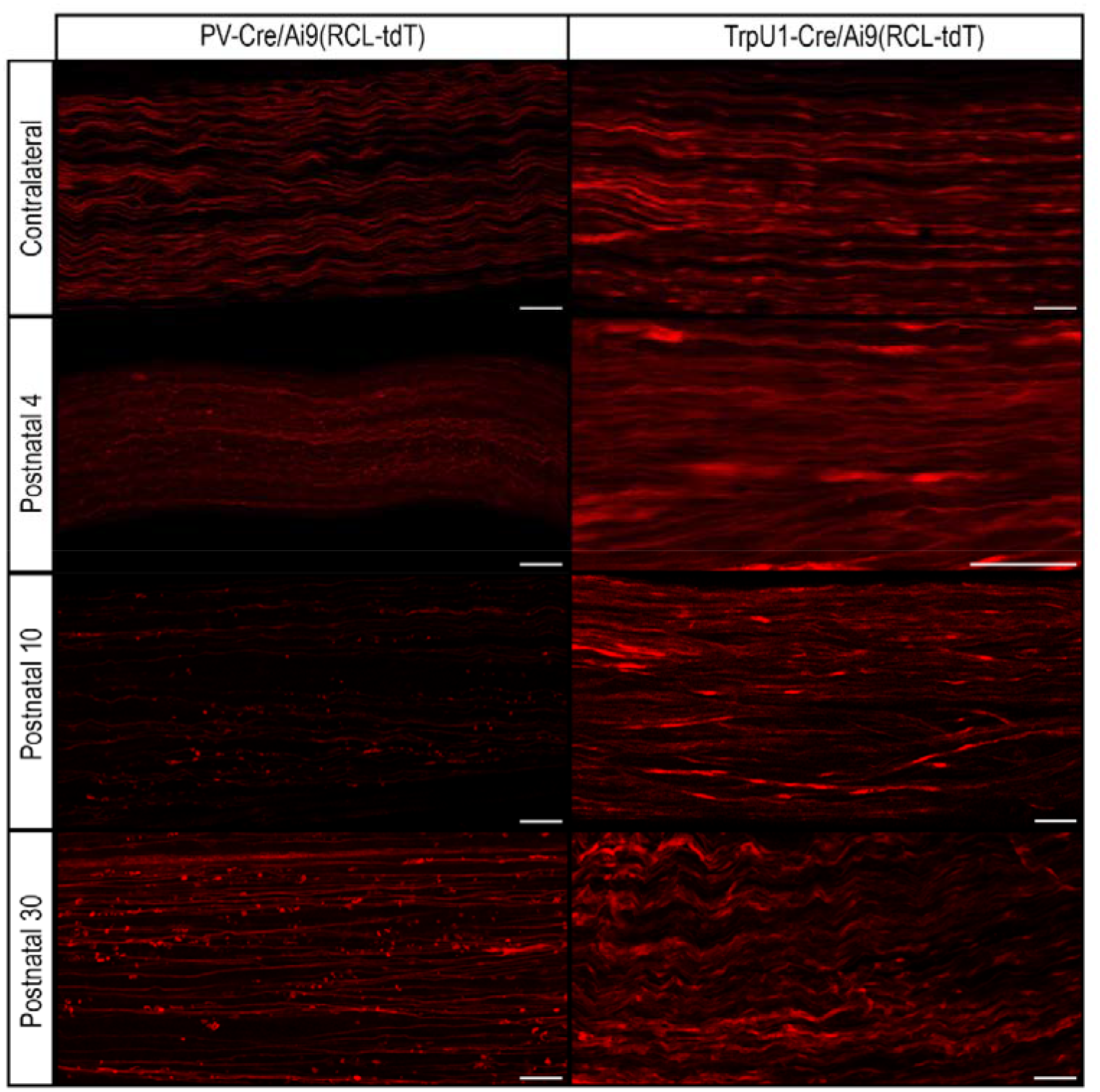
Representative images of longitudinal sections of contralateral and ipsilateral sciatic nerves taken 5 mm distal to the lesion site. Nerves from P4, P10 and P30 PV-Cre/Ai9(RCL-tdT) and TrpV1-Cre/Ai9(RCL-tdT) mice at 7 dpi are shown. tdTomato fluorescence highlights proprioceptive (PV^+^) and nociceptive (TrpV1^+^) axons, allowing visualisation of axonal distribution and continuity across postnatal stages following sciatic nerve crush. Scale bar: 50 µm.

**Figure 4.**
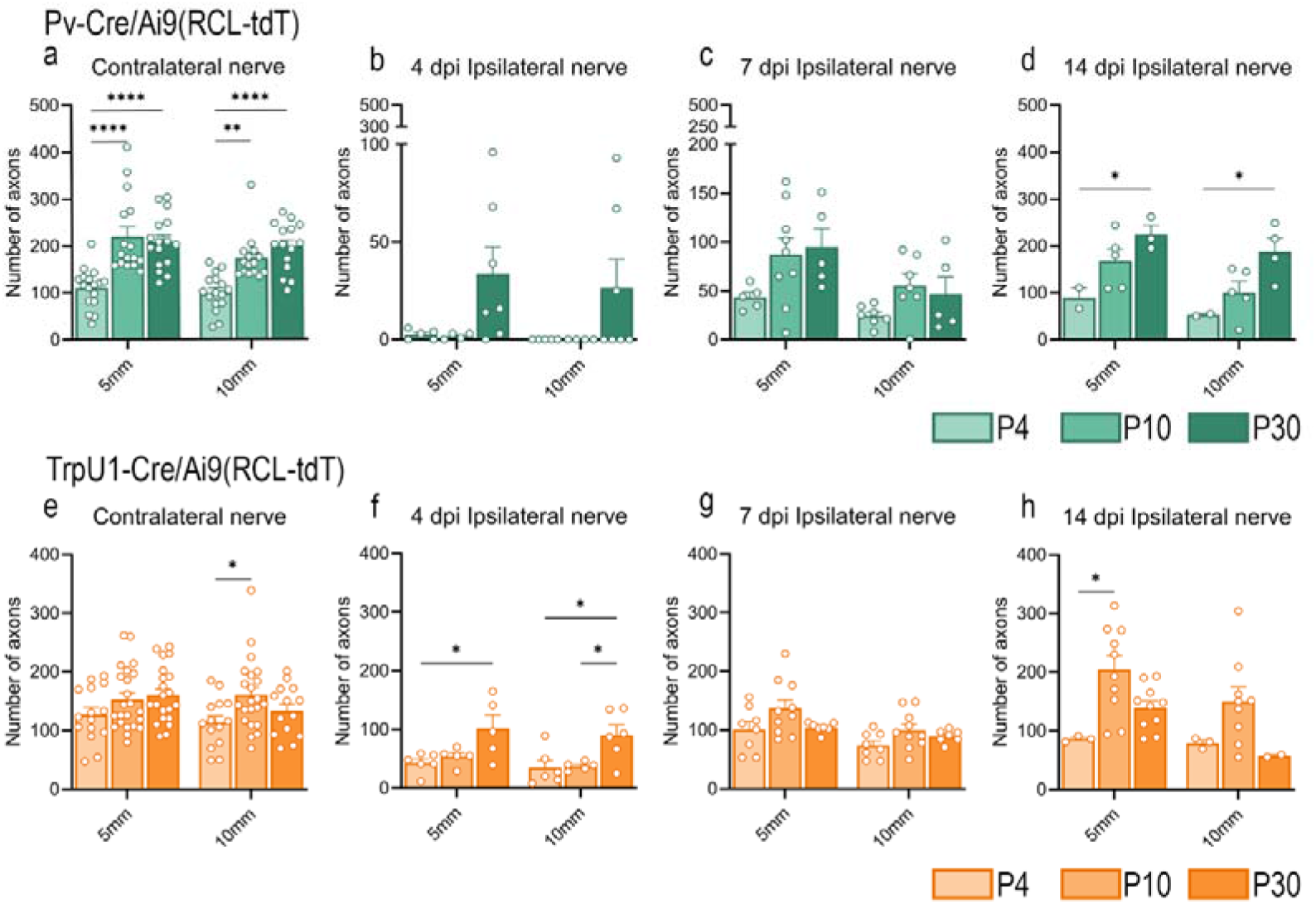
Total number of intact axons counted in contralateral (a, e) and ipsilateral sciatic nerves of P4, P10 and P30 PV-Cre/Ai9(RCL-tdT) and TrpV1-Cre/Ai9(RCL-tdT) mice at 4 (b, f), 7 (c, g) and 14 dpi (d, h). Statistical comparisons were performed using two-way ANOVA followed by Sidak’s post hoc correction. Significance levels: *p<0.05; **p<0.01; ****p<0.0001.

Following a crush injury at different postnatal stages, we quantified the number of proprioceptive and nociceptive axons 5 and 10 mm distal to the lesion site at 4 (Figures 4b, f), 7 (Figures 4c, g) and 14 dpi (Figures 4d, h). As expected, all experimental groups displayed a reduction in axon number 5 and 10 mm distal to the injury at 4 dpi, compared to intact nerves, followed by a progressive increase at 7 and 14 dpi.

To assess the proportion of axons that have regenerated over time across the different postnatal stages, we compared the total number of axons at 4, 7 and 14 dpi with the corresponding contralateral nerve. In proprioceptive neurons (Figures 5a–c), all postnatal stages showed a low percentage of regenerated axons at 4 dpi, which increased at 7 and 14 dpi. Only P30 mice achieved complete regeneration, reaching 100% of contralateral axon numbers at 14 dpi. In nociceptive neurons (Figures 5d– f), all ages showed an approximately 50% reduction in regenerated axons at 4 dpi, followed by an increase at 7 dpi. By 14 dpi, all groups reached 100% 5 mm from the crush site regeneration relative to the contralateral nerve.

**Figure 5.**
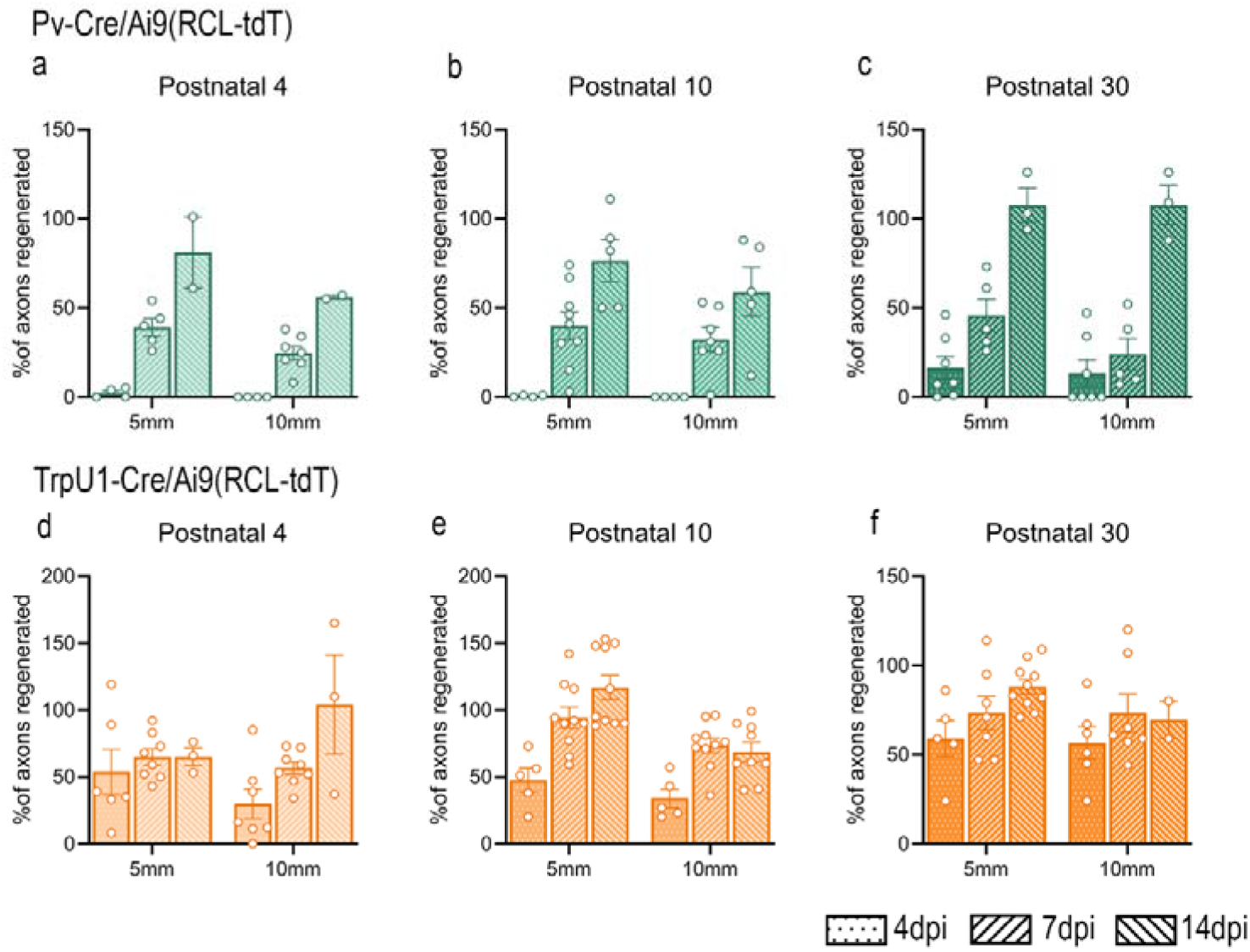
Percentage of regenerated proprioceptive (PV^+^) and nociceptive (TrpV1^+^) axons at 4, 7 and 14 dpi in P4 (a, d), P10 (b, e) and P30 (c, f) PV-Cre/Ai9(RCL-tdT) and TrpV1-Cre/Ai9(RCL-tdT) mice. Percentages were calculated by comparing the number of axons 5 and 10 mm from the crush injury site with the corresponding contralateral nerve.

We next compared the percentage of regenerated axons between P4, P10 and P30 mice at 5 and 10 mm distal to the lesion site (Figure 6). For proprioceptive neurons (Figures 6a, b), P30 mice reached full regeneration (100%) at 10 mm by 14 dpi, statistically significant compared with both P4 and P10 mice. For nociceptive neurons (Figures 6c, d), at 7 dpi P10 mice displayed a higher percentage of regenerated axons than P4 mice (p<0.05), whereas at 14 dpi, this difference was significant compared to both P4 and P30 mice. However, when axons were quantified 10 mm from the injury site, no differences were observed between postnatal stages.

**Figure 6.**
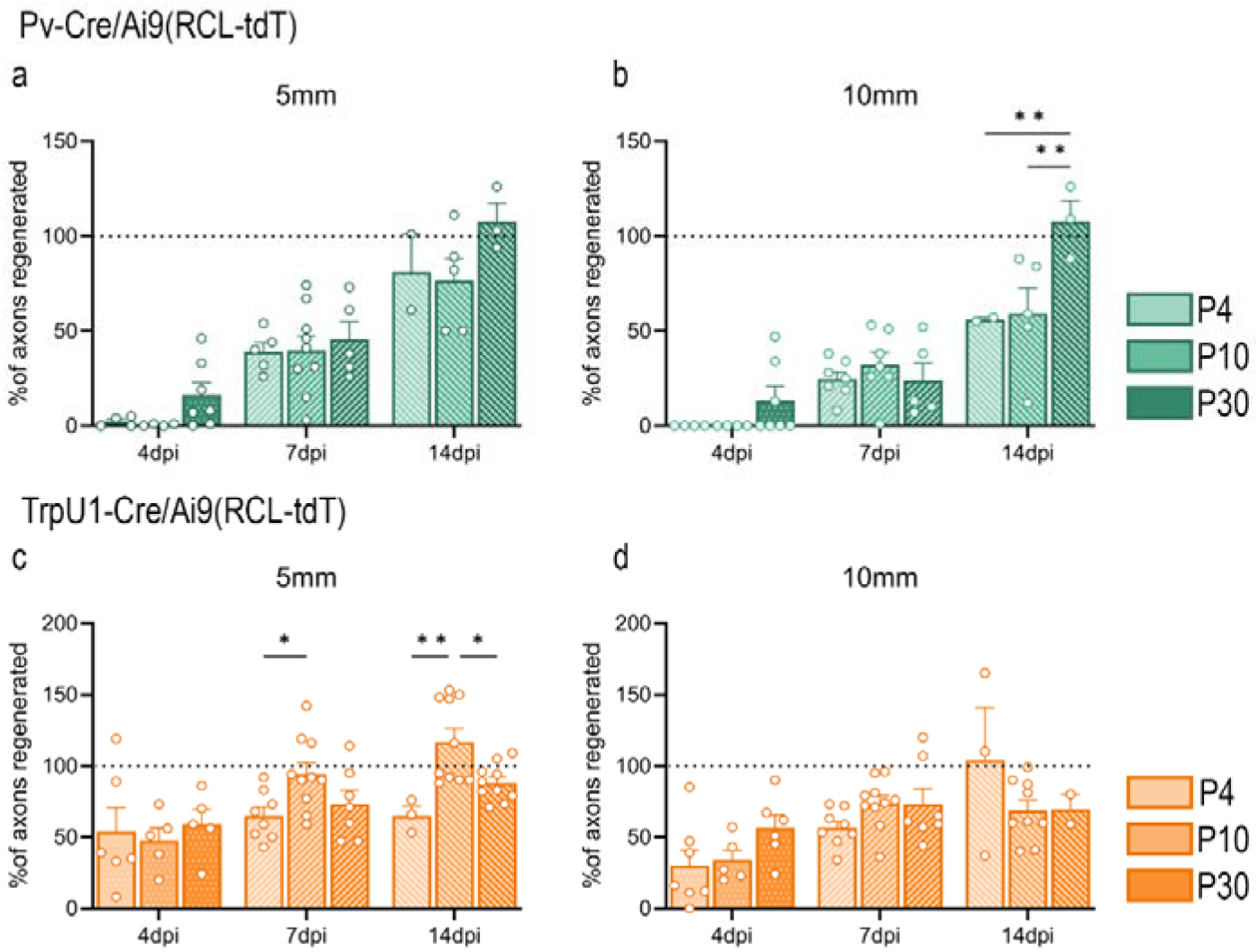
Comparison of the percentage of regenerated proprioceptive (PV^+^) and nociceptive (TrpV1^+^) axons at 5 mm (a, c) and 10 mm (b, d) distal to the lesion site in P4, P10 and P30 mice. Statistical analyses were performed using two-way ANOVA with Sidak’s post hoc test. Significance levels: *p<0.05; **p<0.01.

### Transcriptomic analyses from injured DRGs during the postnatal stages

To investigate the transcriptomic differences underlying regeneration at different postnatal stages, we performed RNAseq on L3-L4 DRGs from injured P4, P10, and P30 mice. DRGs from age-matched uninjured mice (P4+7, P10+7 and P30+7) served as controls. The dataset included 4-5 samples per group, with sequencing yielding an average of ∼23.5 million paired end reads per sample (Figure 7a). PV4CR2I, PV4CR7I and PV4CT2I samples were removed from the analysis due to a number of clean reads inferior to 10.000.000 (Figure 7a, bars in red). UMAP analysis (Figure 7b) shows clustering of the samples based on their gene expression profiles.

**Figure 7.**
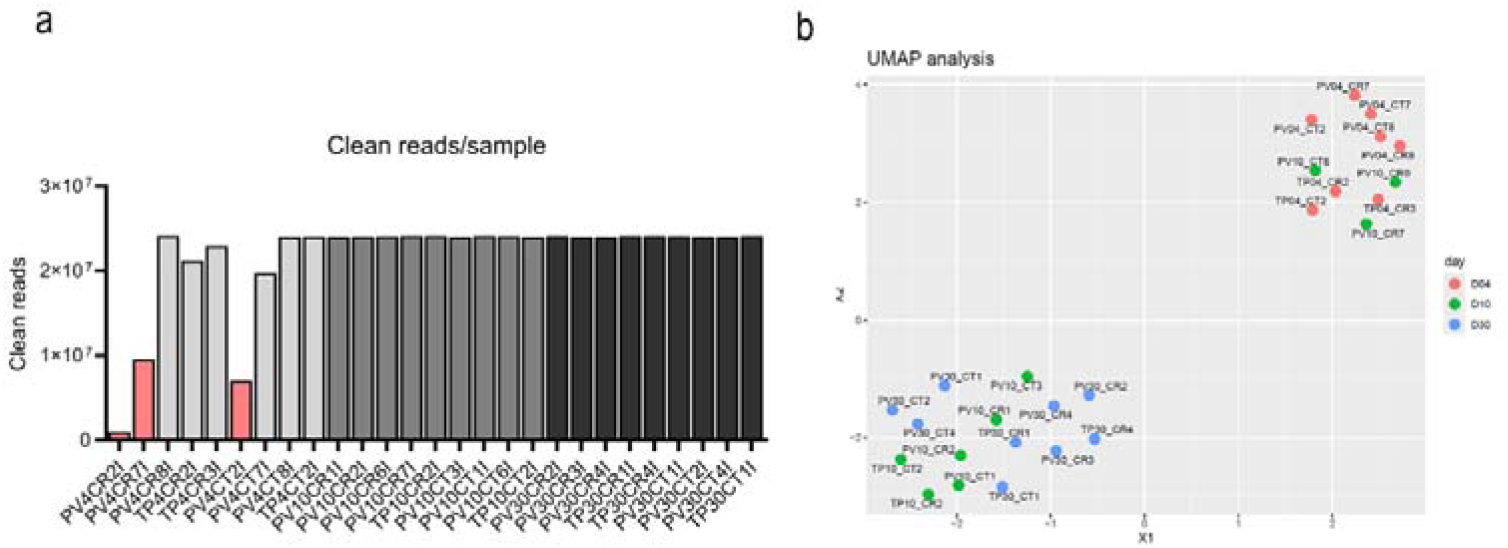
Quality control and exploratory analysis of the transcriptomic dataset. (a) Distribution of the clean reads per sample after a quality filter, showing an homogenous coverage among 27 samples analysed. (b) UMAP analysis of the global clustering of DRGs samples, D04, D10 and D30. Each dot represents an individual sample, showing spatial segregation by condition and time.

#### Gene Set Enrichment Analysis (GSEA)

To investigate whether sets of genes exhibited a consistent increase or decrease in expression across different postnatal stages following injury, we performed gene set enrichment analysis (GSEA). Comparisons were made between the injured group and the corresponding age-matched control group (Injured P4 vs. Intact P4+7, Injured P10 vs. Intact P10+7, Injured P30 vs. Intact P30+7). Using our custom list of RAGs and RStudio software (RStudio Team, 2024, version 2024.12.0+467), we conducted GSEA (Figure 8a-c). We subsequently focused on the leading-edge RAGs identified via GSEA to pinpoint the genes most significantly enriched at each postnatal stage. In P30 animals, these classical RAGs, were upregulated compared with their noninjured counterparts (Figure 8a). In contrast, compared with noninjured animals of the same age, injured P4 and P10 mice did not exhibit any significantly upregulated RAGs (Figure 8b, c). Following the GSEA findings, we next examined the expression levels of the identified leading-edge RAGs by assessing their transcripts per million (TPM) values across the experimental groups (Figure 9a-f). Consistent with the GSEA results, injured P30 animals presented visibly greater expression of these RAGs than did their intact P30+7 counterparts. In contrast, the TPM values for these RAGs in both injured P10 and injured P4 animals showed minimal discernible changes relative to their respective age-matched intact control groups.

**Figure 8.**
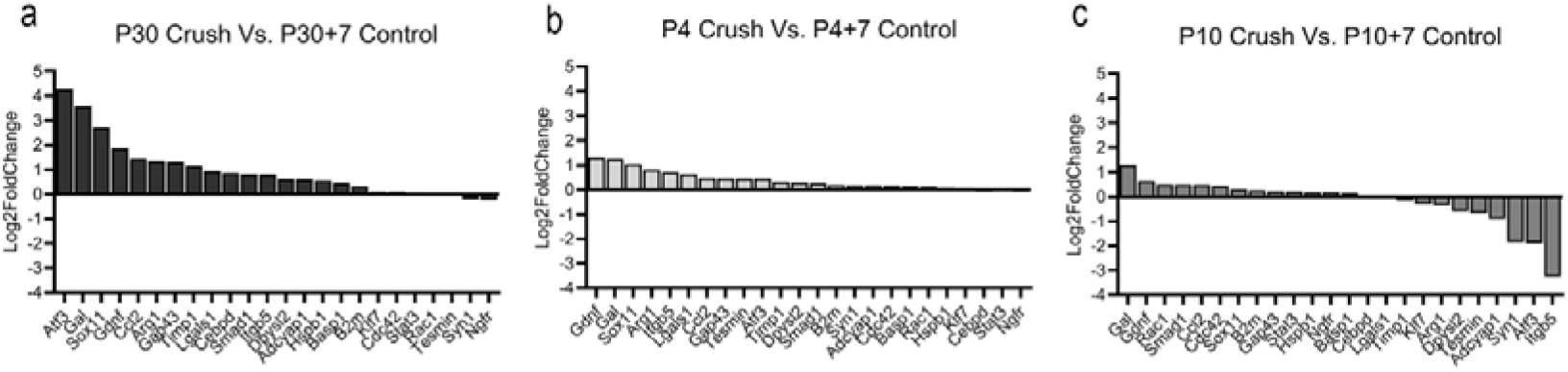
GSEA showing Log2FoldChange of RAGs (Atf3, Gal, Sox11, Gdnf, Ccl2, Arg1, Gap43, Timp1, Lgals1, Cebpd, Smad1, Itgb5, Dpysl2, Adycap1, Hspb1, Basp1, B2m, Klf7, Cdc42, Stat3, Rac1, Tesmin, Syn1, Ngfr) across P4, P10 and P30 injured L3-L5 DRGs compared to control ones.

**Figure 9.**
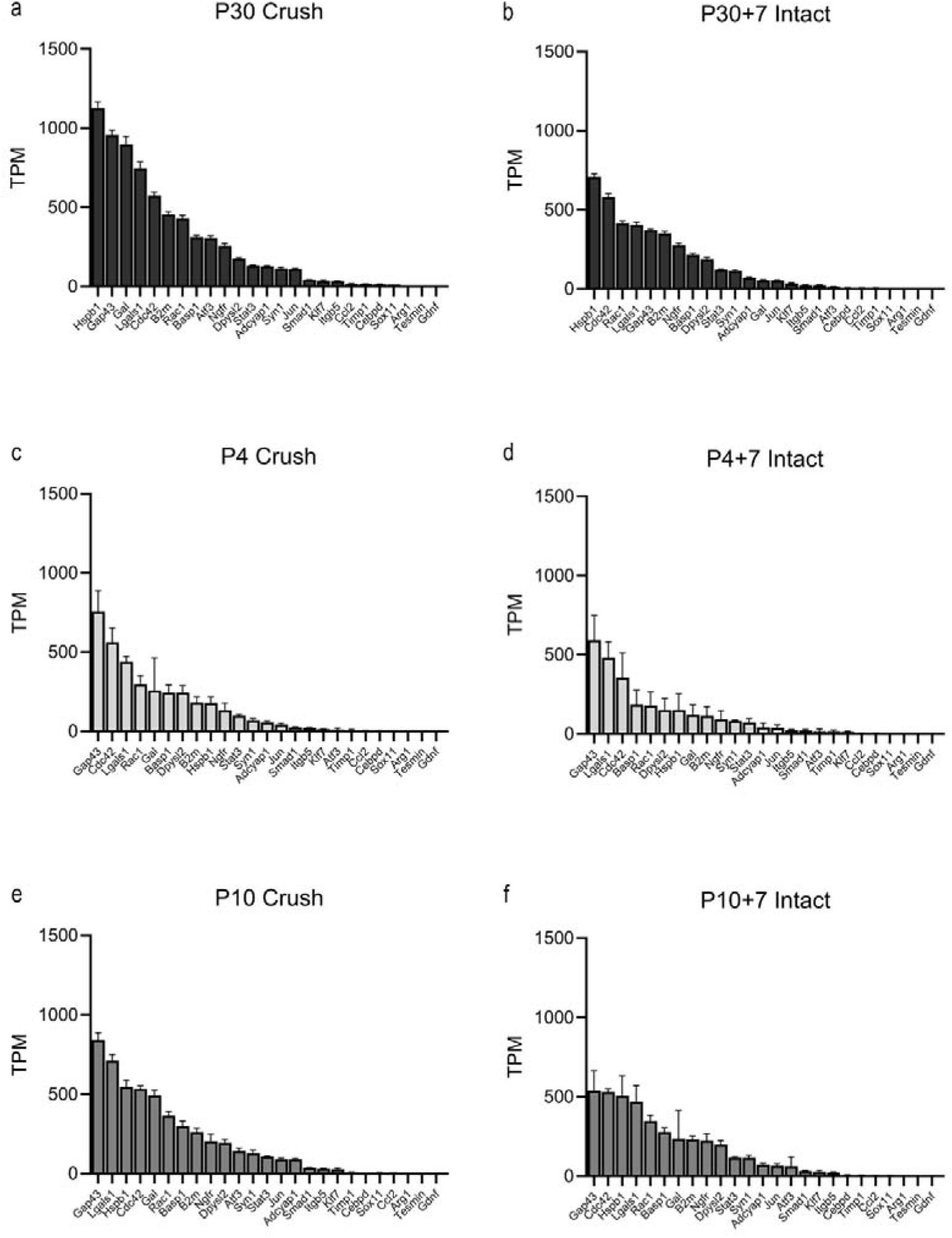
(a–f) Transcripts per million (TPM) of selected RAGs in intact and injured DRGs from P30, P30+4, P4, P4+7, P10 and P10+7 mice. The data are presented as the means ± SEMs (*n* = 3-5 mice per group).

#### Pathway enrichment analysis

To investigate the molecular response of sensory neurons to sciatic nerve crush at different postnatal stages, we performed pathway enrichment analysis (Figure 10, Supplementary Table 1). When the analysis was performed, only the P30 mice exhibited upregulated signalling pathways with a statistical significance of *padj*<0.05, whereas the animals injured at P4 and P10 showed none. Therefore, to gain an approximate view of pathways that might be either up-or downregulated, we selected those displaying a *p*-value<0.05, even though their *padj* exceeded 0.05 (Supplementary Figure 2).

**Figure 10.**
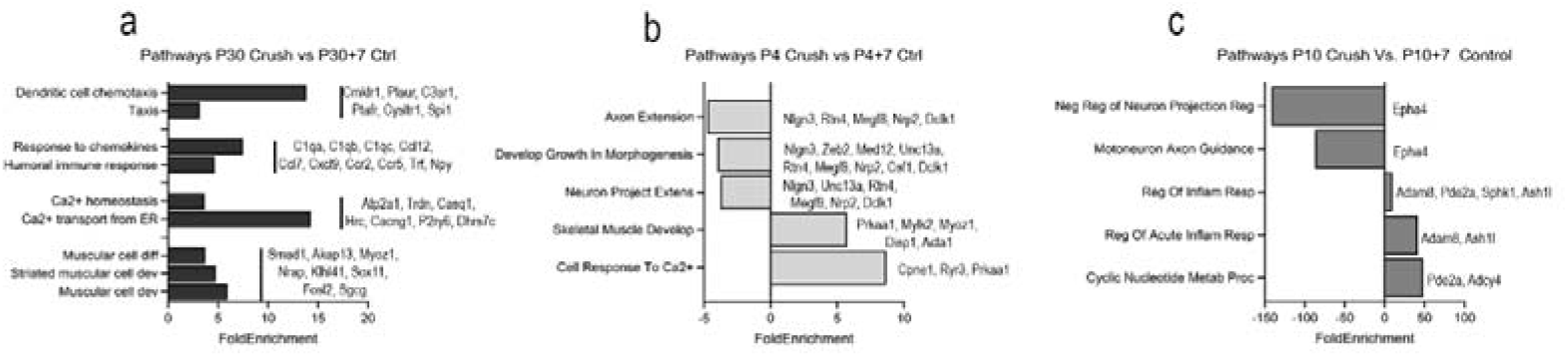
Pathway enrichment analysis. (a) Molecular pathways enriched in P30 crush animals compared with P30+7 control animals. (b) Molecular pathways enriched in P4 crush animals compared with P4+7 control animals. (c) Molecular pathways enriched and downregulated in P10 crush animals compared with those in P10+7 control animals.

The analysis revealed distinct stage-dependent profiles. In injured P30 mice, compared to age-matched controls (Figure 10a), we observed upregulation of pathways related to dendritic cell chemotaxis, response to chemokines, and calcium homeostasis, among others. In injured P4 mice (Figure 10b), most identified pathways—including those involved in axon extension and morphogenetic development—were downregulated; however, pathways associated with the response to calcium were upregulated. In injured P10 mice compared to uninjured age-matched animals (Figure 10c), pathways related to axon extension and neuronal projection were downregulated, whilst pathways associated with the inflammatory response were upregulated.

Based on the enriched signalling pathways (upregulated or downregulated), we selected a series of genes primarily related to the immune response and the recruitment of macrophages and satellite cells (Figure 11a, Table 1), as well as others associated with nervous system development, axonal guidance, and synaptic plasticity (Figure 11b, Table 2).

**Figure 11.**
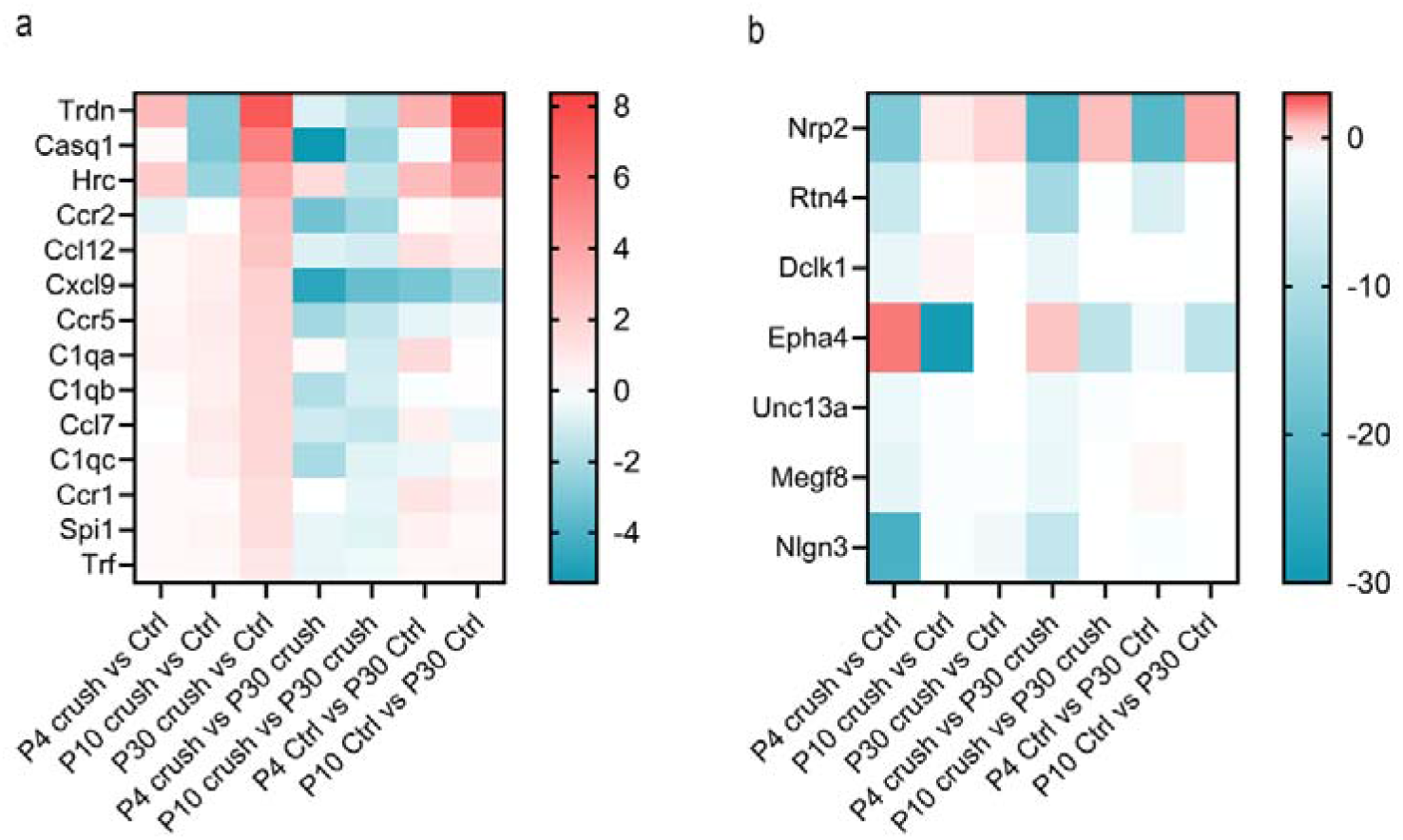
a) Heatmap representing the differential expression (Log2 fold change) of genes related to the inflammatory response across developmental stages (P4, P10, P30) and experimental conditions (crush, control) b) Heatmap representing the differential expression (Log2 fold change) of genes related to axonal growth across developmental stages (P4, P10, P30) and experimental conditions (crush, control). Colour scale indicates Log2 fold change, with red representing upregulation and blue representing downregulation compared to the indicated comparison groups.

**Table 1.** Differential expression of genes related to the inflammatory response across developmental stages and experimental conditions. Log2 fold change (Log2FC) and adjusted p-values (padj) for each gene were calculated for the indicated experimental groups. Comparisons include P4, P10, and P30 after nerve crush versus their respective age matched controls (Ctrl), and pairwise comparisons between ages for both crush and control conditions (P4 vs P30, P10 vs P30). Genes with padj<0.05 are considered significantly differentially expressed.

|  | P4 Crush vs Ctrl |  | P10 Crush vs Ctrl |  | P30 Crush vs Ctrl |  | P4 vs P30 (Crush) |  | P10 vs P30 (Crush) |  | P4 vs P30 (Ctrl) |  | P10 vs P30 (Ctrl) |  |
| --- | --- | --- | --- | --- | --- | --- | --- | --- | --- | --- | --- | --- | --- | --- |
| Gene ID | Log2FC | padj | Log2FC | padj | Log2FC | padj | Log2FC | padj | Log2FC | padj | Log2FC | padj | Log2FC | padj |
| Ccl12 | 0,36 | 0,912 | 0,68 | 1,000 | 2,5 | 2,7E-06 | -0,78 | 0,098 | -1,03 | 0,168 | 1,35 | 0,010 | 0,79 | 0,940 |
| Ccl7 | -0,01 | 0,999 | 0,89 | 1,000 | 1,74 | 2,6E-03 | -1,09 | 0,021 | -1,36 | 0,024 | 0,67 | 0,194 | -0,5 | 1,000 |
| Ccr5 | 0,41 | 0,921 | 0,85 | 1,000 | 1,89 | 6,2E-03 | -2,09 | 1,8E-03 | -1,36 | 0,082 | -0,6 | 0,340 | -0,32 | 1,000 |
| Ccr2 | -0,64 | 0,944 | 0,07 | 1,000 | 2,74 | 3,4E-02 | -3,22 | 0,002 | -2,13 | 0,109 | 0,16 | 0,895 | 0,54 | 1,000 |
| Cxcl9 | 0,32 | 0,987 | 0,73 | 1,000 | 2 | 4,1E-02 | -4,75 | 8,3E-08 | -3,45 | 0,000 | -3,07 | 0,002 | -2,18 | 0,101 |
| Ccr1 | 0,27 | 0,946 | 0,23 | 1,000 | 1,46 | 4,3E-02 | -0,02 | 0,971 | -0,57 | 0,941 | 1,16 | 0,029 | 0,65 | 1,000 |
| C1qa | 0,55 | 0,867 | 0,72 | 1,000 | 1,87 | 2,2E-02 | 0,27 | 0,689 | -1,04 | 0,502 | 1,59 | 0,010 | 0,11 | 1,000 |
| C1qb | 0,14 | 0,962 | 0,68 | 1,000 | 1,77 | 2,9E-07 | -1,77 | 2,2E-08 | -0,96 | 0,021 | -0,14 | 0,704 | 0,13 | 1,000 |
| C1qc | 0,24 | 0,923 | 0,76 | 1,000 | 1,69 | 9,6E-06 | -1,94 | 1e-08 | -0,72 | 0,248 | -0,49 | 0,189 | 0,21 | 1,000 |
| Trf | 0,25 | 0,885 | 0,34 | 1,000 | 1,08 | 3,3E-03 | -0,51 | 0,092 | -0,39 | 0,836 | 0,32 | 0,318 | 0,35 | 1,000 |
| Spi1 | 0,28 | 0,947 | 0,44 | 1,000 | 1,44 | 5,0E-02 | -0,52 | 0,325 | -0,71 | 0,732 | 0,64 | 0,250 | 0,29 | 1,000 |

**Table 2.** Differential expression of genes related to axonal growth across developmental stages and experimental conditions. Log2 fold change (Log2FC) and adjusted p-values (padj) for each gene related to axonal growth were calculated for the indicated experimental groups. Comparisons include P4, P10, and P30 after nerve crush versus their respective controls (Ctrl), and pairwise comparisons between ages for both crush and control conditions (P4 vs P30, P10 vs P30). Genes with padj<0.05 are considered significantly differentially expressed.

|  | P4 Crush vs Ctrl |  | P10 Crush vs Ctrl |  | P30 Crush vs Ctrl |  | P4 vs P30 (Crush) |  | P10 vs P30 (Crush) |  | P4 vs P30 (Ctrl) |  | P10 vs P30 (Ctrl) |  |
| --- | --- | --- | --- | --- | --- | --- | --- | --- | --- | --- | --- | --- | --- | --- |
| Gene | Log2FC | padj | Log2FC | padj | Log2FC | padj | Log2FC | padj | Log2FC | padj | Log2FC | padj | Log2FC | padj |
| Nlgn3 | -22,98 | 4,1E-11 | -7,39 | 0,030 | -0,87 | 8,2E-01 | -0,85 | 1,000 | -0,14 | 1,000 | -0,27 | 1,000 | -1,96 | 1,000 |
| Unc13a | -2,48 | 9,7E-04 | -2,44 | 4,0E-05 | -0,11 | 8,6E-01 | -0,62 | 1,000 | -0,68 | 0,832 | -0,21 | 1,000 | -0,16 | 1,000 |
| Rtn4 | -6,67 | 1,7E-03 | -11,48 | 01,3E-12 | -4,76 | 2,4E-04 | -0,06 | 1,000 | -0,4 | 1,000 | -0,29 | 1,000 | 0,05 | 1,000 |
| Megf8 | -3,26 | 1,1E-02 | -2,5 | 0,007 | 0,13 | 9,0E-01 | -0,74 | 1,000 | -0,32 | 1,000 | -0,21 | 1,000 | -0,63 | 1,000 |
| Nrp2 | -15,93 | 1,8E-02 | -21,83 | 01,7E-06 | -21,07 | 9,9E-06 | 0,32 | 1,000 | 1,01 | 1,000 | 1,4 | 1,000 | 0,66 | 1,000 |
| Dclk1 | -2,8 | 4,8E-02 | -2,83 | 0,002 | -0,05 | 9,6E-01 | 0,19 | 1,000 | -0,15 | 1,000 | -0,36 | 1,000 | -0,02 | 1,000 |
| Epha4 | 2,1 | 9,4E-01 | 0,91 | 0,825 | -1,33 | 7,5E-01 | -29,13 | 1,7E-14 | -8,2 | 0,187 | -8,29 | 0,368 | -0,08 | 1,000 |

When examining genes related to the immune response (Table 1) (Ccl12, Ccl7, Ccr5, Ccr2, Cxcl9, Ccr1, C1qa, C1qb, C1qc, Trf, and Spi1), we observed significant overexpression in injured animals at P30 compared with uninjured P30+7 animals (Log2FC>1, padj<0.05). This pattern was not observed in injured animals at P4 or P10 when compared with their respective age-matched controls (padj>0.05). When comparing these same genes between injured animals at P4 or P10 versus injured animals at P30, several genes were found to be downregulated at P4 (Ccl7, Ccr5, Ccr2, Cxcl9, C1qb, C1qc) and at P10 (Ccl7, Cxcl9, C1qb) (Log2FC<0, padj<0.05). However, when comparing control animals at P4 versus control animals at P30, certain genes showed higher expression levels (Ccl12, Ccr1, C1qa; Log2FC>1, padj<0.05), whilst others exhibited lower expression (Cxcl9; Log2FC<0, padj<0.05).

When examining genes related to nervous system development, axonal guidance, and synaptic function (Table 2) (Nlgn3, Unc13a, Rtn4, Megf8, Nrp2, Dclk1, Epha4), comparison of injured animals at P4 and P10 with their respective age-matched controls revealed that all genes except Epha4 exhibited lower expression levels (Log2FC<0; padj<0.05). This pattern was not observed in injured animals at P30 (except for the Nrp2 gene). When comparing injured animals at P4 and P10 with injured animals at P30, only the Epha4 gene showed lower expression in injured animals at P4 (Log2FC<0; padj<0.05). No differences were observed between control animals at P4 and P10 versus control animals at P30 (Log2FC≈0; padj>0.05).

### Neurite outgrowth after a conditioning lesion

To assess the intrinsic capacity for neurite outgrowth at different postnatal stages, and to determine whether a peripheral nerve injury is conditioning sensory neurons to enhance this outgrowth, we quantified neurite extension under basal conditions (P4, P10, and P30, Figure 12) and 7 days after a conditioning lesion (Figure 13). Immunostaining for β-III-tubulin revealed differences in neuronal morphology and neurite length between ages and experimental conditions (Figure 12, 13).

**Figure 12.**
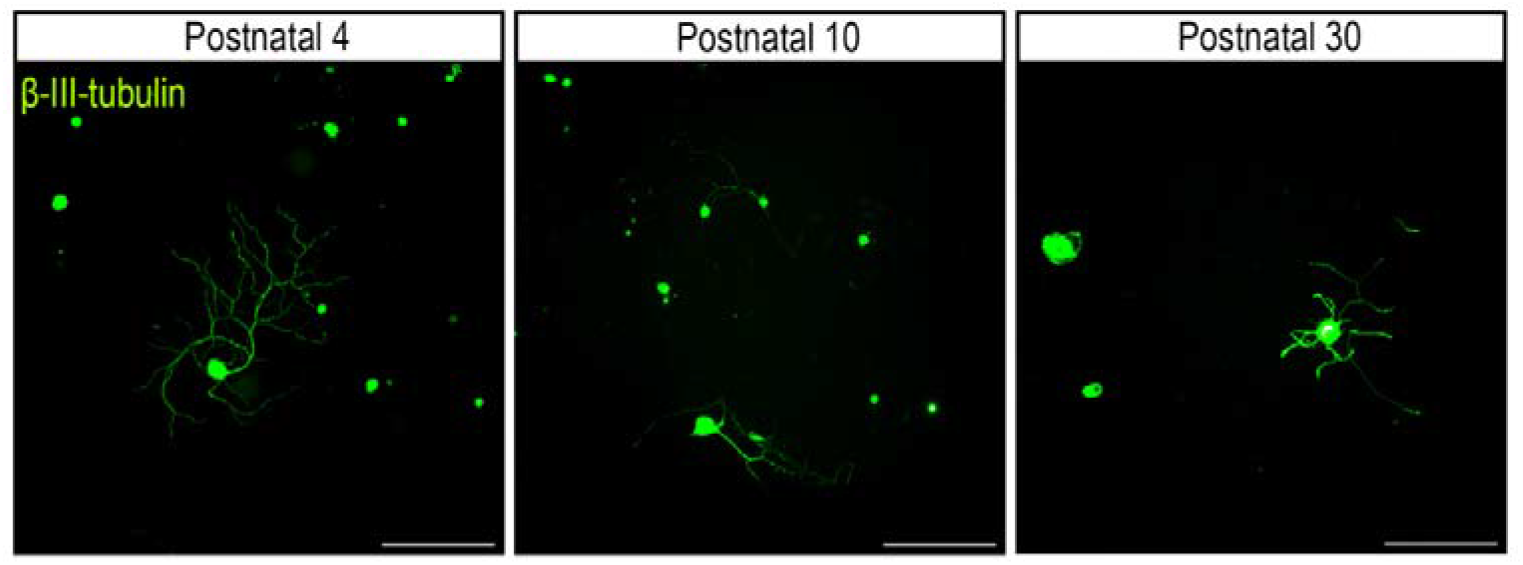
Representative images of β-III-tubulin immunostained primary sensory neurons from lumbar DRG of P4, P10 and P30 mice (left to right), cultured for 24h. Scale bar: 100 µm.

**Figure 13.**
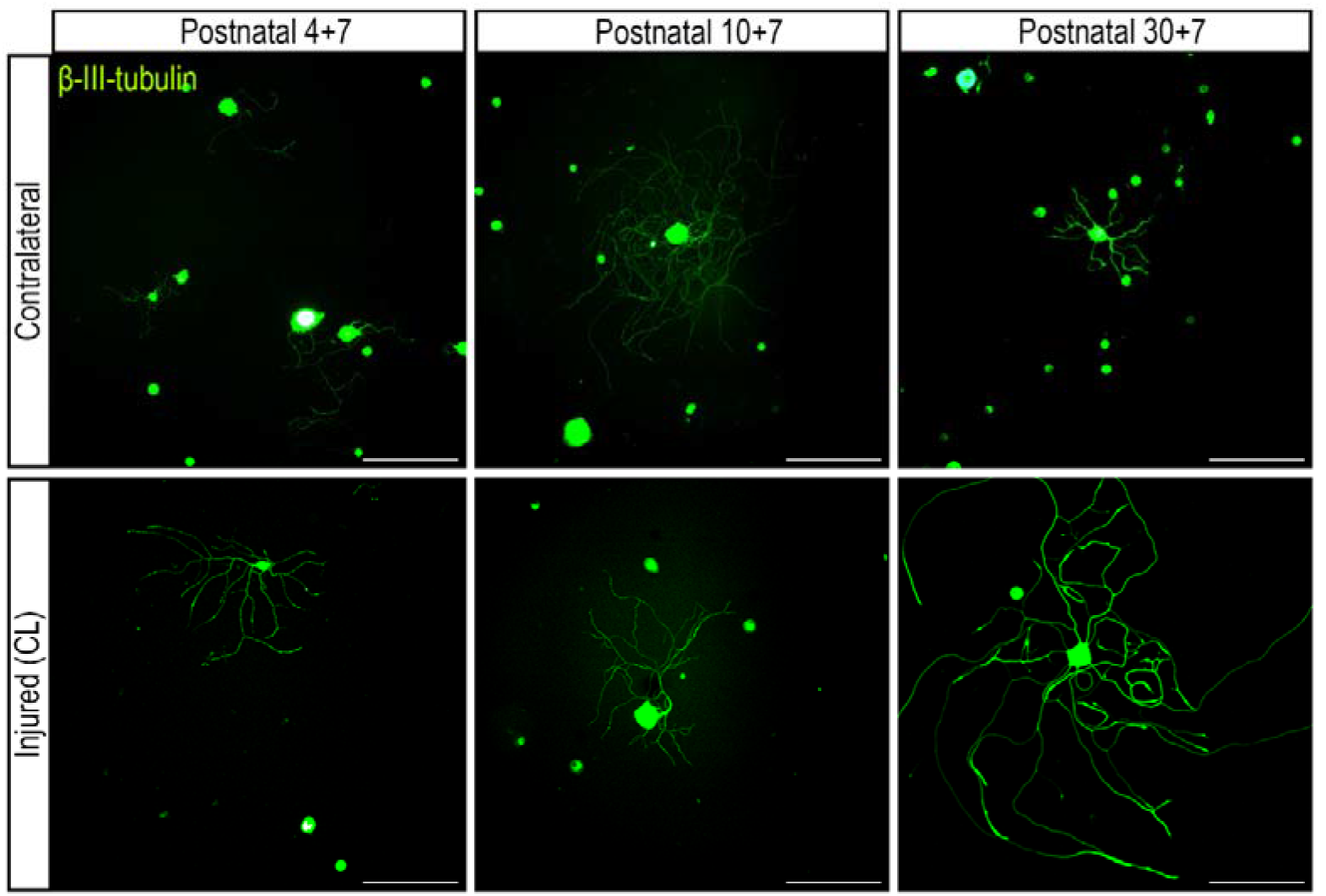
Representative images of cultured primary sensory neurons from lumbar DRG of P4, P10 and P30 mice (left to right) with a prior *in vivo* crush injury to the right sciatic nerve (conditioning lesion) 7 days before (P4+7, P10+7, P30+7). Contralateral (upper row), and injured-ipsilateral-(conditioning lesion, lower row) conditions are shown. Neurons were immunostained with the panmarker β-III-tubulin in. Scale bar: 100 µm

Under basal (uninjured) conditions sensory neurons from P4, P10, and P30 cultured for 24h showed a similar morphology and heterogeneity, with neurons without neurites and others extending neurites of different lengths (Figure 14). There were no statistically significant differences in the percentage of neurons bearing neurites (Figure 14a), mean neurite length (Figure 14b), or the length of the longest neurite (Figure 13c) between these ages, although all parameters had the tendency to be higher in P4 neurons.

**Figure 14.**
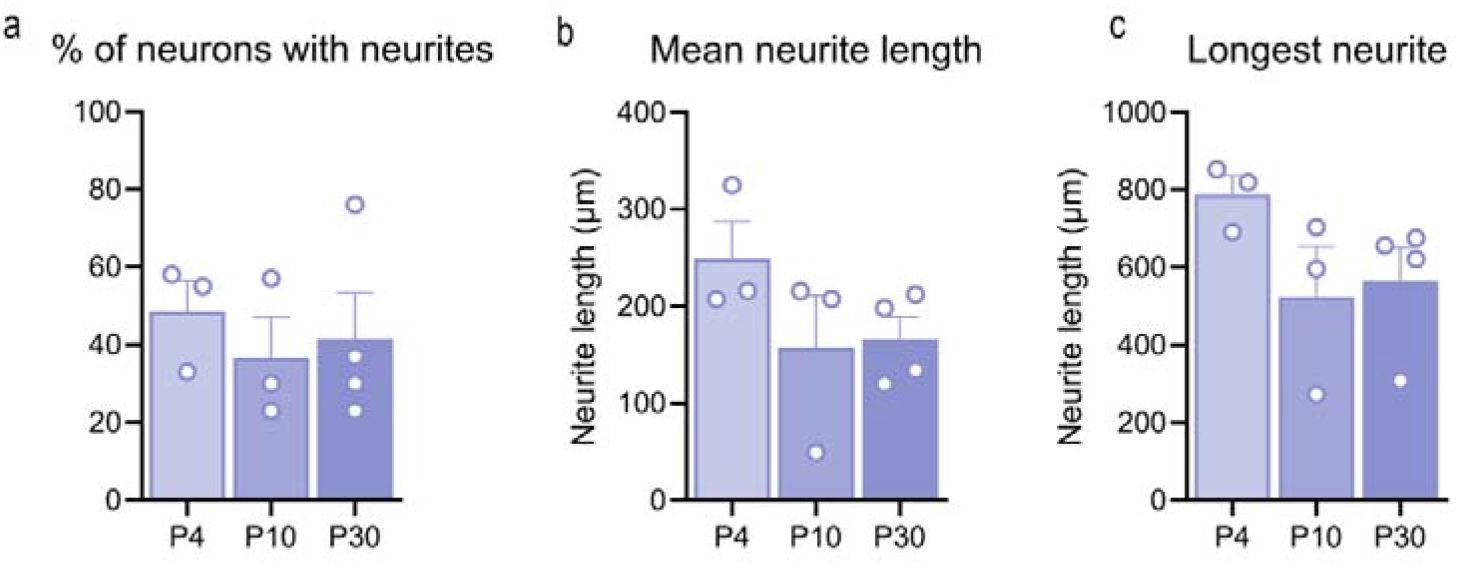
Neurite outgrowth of neurons at different postnatal stages. (a) Percentage of neurons with neurites at P4, P10 and P30. (b) Mean neurite length of neurons with neurites at P4, P10 and P30. (c) Longest neurite at P4, P10 and P30. Statistical analyses were performed using One-way ANOVA with Tukey post hoc test.

We also evaluated the effect of the conditioning lesion at different postnatal stages by focusing on the longest neurite for each culture. We observed a marked increase in that length following conditioning lesion at P30+7 compared to its contralateral side (p<0.05). In contrast, P4+7 mice displayed no difference between sides (p>0.05), and P10+7 mice showed a reduction in the longest neurite after injury relative to their contralateral side (p<0.05) (Figure 15a-c).

**Figure 15.**
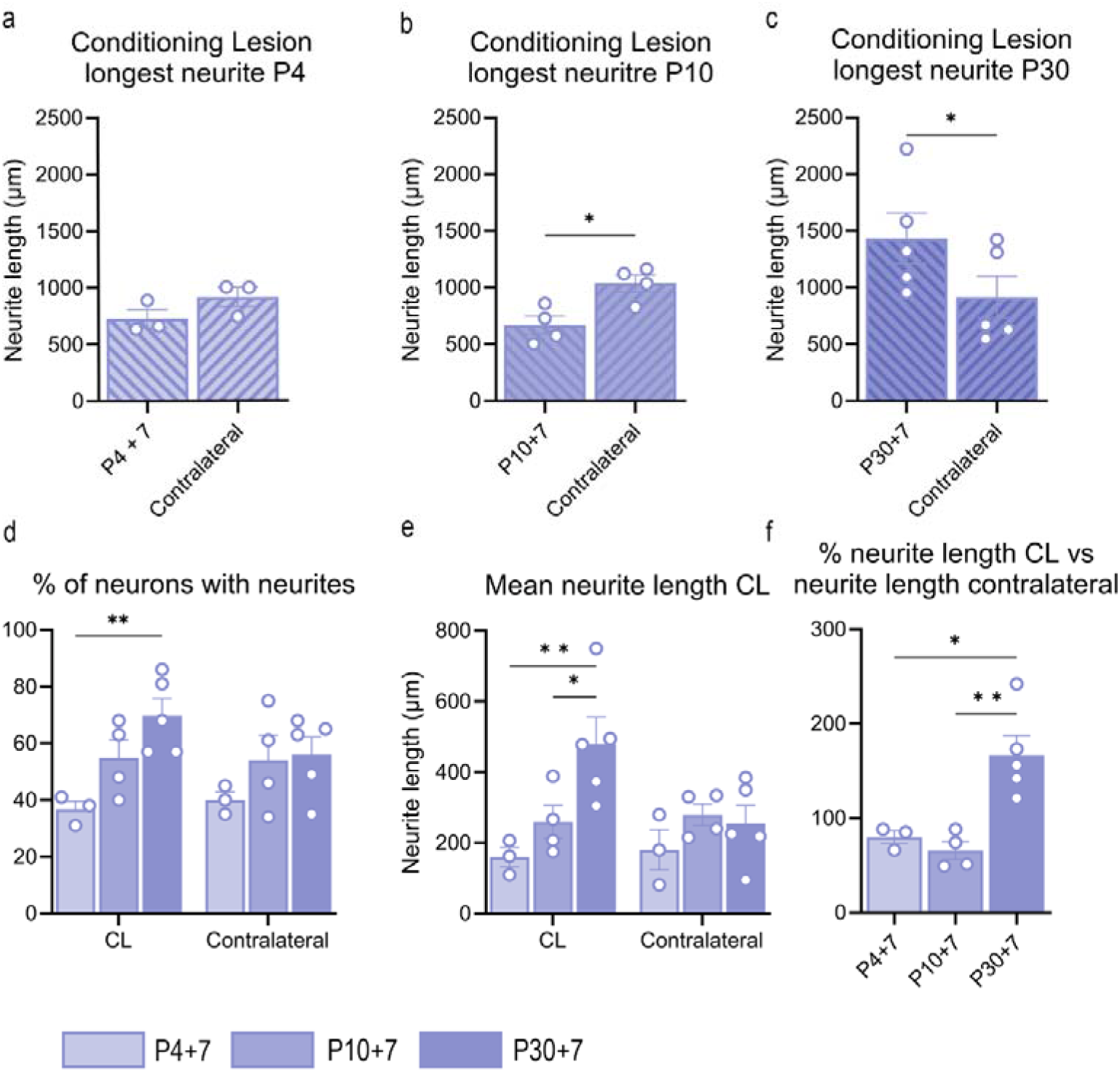
Quantitative analysis of neurite outgrowth following a conditioning lesion at different postnatal stages. (a-c) Longest neurite length for neurons at P4+7 (d), P10+7 (e), and P30+7 (f) in the CL and contralateral groups. Statistical analyses were performed using a paired t test *p < 0.05, **p < 0.01. (d) Percentage of neurons with neurites 7 dpi after a conditioning lesion in the ipsilateral injured side and the contralateral side at P4, P10 and P30. Statistical analyses were performed using two-way ANOVA with Sidak’s post hoc test. Significance levels: **p<0.01 (e) Mean neurite length 7 dpi after a conditioning lesion in the ipsilateral injured side and the contralateral side at P4, P10 and P30. Statistical analyses were performed using two-way ANOVA with Sidak’s post hoc test. Significance levels: *p<0.05 (f) Comparison of longest neurite length between CL and contralateral groups at P4+7, P10+7 and P30+7. Statistical analyses were performed using One-way ANOVA with Tukey post hoc test. Significance levels: *p<0.05 ** p<0.01.

When comparing other parameters between ages, we observed that a conditioning lesion increases the percentage of neurons with neurites at P30+7 compared to P4+7 mice; this effect was not observed on the contralateral side (Figure 15d). Comparisons of mean neurite length revealed that P30+7 mice exhibited significantly greater neurite length than P4+7 (p<0.01) and P10+7 (p<0.05) mice on the ipsilateral side, but not on the contralateral side (Figure 15e). Interestingly, neurons from the contralateral side behave similar to basal neurons of the same age (P4+7 vs P10 and P30+7 vs P30), although with a tendency to show higher values in all the parameters evaluated (Supplementary Figure 2).

For a fair comparison of the effects of the CL among postnatal stages, we calculated the percentage of change in the length of the longest neurite in axotomized neurons compared to contralateral non injured ones. We observed a 66% increase in the length of the longest neurite following injury in P30 mice. In contrast, injury at P4 and P10 resulted in a decrease of approximately 20% and 25%, respectively. Overall, these results confirm that a conditioning lesion at P30 significantly enhances neurite outgrowth compared to P4 (p<0.05) and P10 (p<0.01) (Figure 15f)

## Discussion

In this study, our main objective is to explore if genetic program activated by injury in primary sensory neurons differ at different postnatal stages. To achieve this, we integrate morphological, molecular, transcriptomic, and *in vitro* approaches, aiming to shed light on the fundamental mechanisms that drive the regenerative response in the postnatal peripheral nervous system.

To evaluate the regenerative capacity of sensory neurons, we selected two paradigmatic subtypes, proprioceptors and nociceptors. Interestingly, they present differences in the regenerative response after a peripheral nerve injury in adult stages, although share common pro-regenerative pathways, too (Bolívar et al., 2024).

We observe a notable absence of death of both types of neurons during early postnatal development (P4 and P10) following peripheral nerve injury, which contrasts with the marked reduction of motoneurons reported in our previous work (Molina-Esteve et al., 2025). In fact, massive loss of postnatal injured motoneurons have been extensively documented in the literature (Lowrie, Krishnan, and Vrbova 1982; Liang Low et al., 2003; Greensmith, Hasan, and Vrbová 1994; Lowrie and Vrbová & 1992) and it is restricted to early postnatal stages, whereas loss of sensory neurons is more moderate (Bahadori et al., 2001) and can be extended to juvenile (Kemp et al., 2015a) and even adults (Cooper et al., 2024; Tandrup et al., 2000) where it has been described that the small non-peptidergic nociceptors are most susceptible to axotomy. If we assume that the same population of neurons would die at early postnatal stages, only populations including these small neurons should be vulnerable to injury. Therefore, no reduction in proprioceptors but in nociceptors would be expected. Unfortunately, the analysis in the whole DRG stacked in the confocal microscope could limited our capacity to detect the whole population of TrpV1^+^ neurons, in a prominent manner the smaller ones, and therefore, prevented us to detect its loss.

Both proprioceptors and nociceptors exhibited a high capacity for regeneration following sciatic nerve injury across all postnatal stages (P4, P10, P30), with nociceptive neurons showing significantly greater regeneration than proprioceptive neurons in the early postnatal stages. These findings are consistent with established literature demonstrating that unmyelinated axons possess superior regenerative velocity compared to their myelinated counterparts (Navarro et al., 1994; Bolívar et al., 2024).

Nociceptors exhibited a distinctive regenerative dynamic, and the number of axons measured at 14 dpi reached basal values at all ages at some distances. In fact, more regenerative axons were observed at some points and ages compared to control values. It is well known that the tip of the regenerating axons can extend a bundle of sprouts at early stages of regeneration (Witzel et al., 2005) and therefore, the estimated number of axons can also reflect this phenomenon.

When focusing on proprioceptive neurons, significant differences emerged between the early postnatal groups (P4 and P10) with lower regenerative outcomes than P30 injured animals. These findings parallel our observations with motoneurons described previously (Molina-Esteve et al., 2025), demonstrating superior regenerative capacity in P30 mice attributed to the robust activation of RAGs, that was not observed in P4 and P10 mice.

Similarly, in sensory neurons no overexpression of RAGs was detected after injury during the early postnatal stages, while such overexpression occurred in injured mice at P30. Transcriptomic analyses revealed that classical pro-regenerative pathways were activated only in young adults, whereas pathways related to development and inflammatory processes were regulated in early postnatal mice. By applying a conditioning injury, we could further explore the impact of these different genetic programs in neurite growth. Despite neurons of different ages extend similar neurites in vitro, the conditioning lesion paradigm was only observed in young adult (P30) mice, where the additional injury clearly promoted enhanced growth capacity; this conditioning lesion effect has been extensively reported in the literature (Smith and Skene 1997; Lankford et al., 1998; Dubový et al., 2019; Seo et al.m 2025). In contrast, when applying injuries to P4 and P10 mice, this effect was not observed, supporting the idea that these neurons are in a basal state that inherently promotes growth and therefore, no genetic switch towards a pro-regenerative state is needed for a successful regenerative response. In fact, in P10 animals we paradoxically observed impaired regenerative responses following a conditioning lesion. Therefore, the age-dependent conditioning response observed in this study underscores the fundamental shift from an axonal growth program in neonatal neurons to a neurotransmitter state in mature ones, that need a genetic shift to reactivate their growth capacities.

Our findings provide compelling evidence that RAGs are robustly upregulated exclusively at P30, remain dormant at P4, and exhibit active downregulation at P10. Our findings are in line with our results in motoneurons (Molina-Esteve et al., 2025) and with previous works that point that P10 represents a critical developmental window characterized by compromised regenerative capacity of peripheral neurons (Fitzgerald and McKelvey 2016).

It is well established that a dramatic reduction of axon growth rates is observed in neonatal ages, since when developing neurons reach their target, switch from an outgrowth program to a neurotransmitter mode (Goldberg et al., 2002). Peripheral neurons retain the ability to reactivate this growth state after injury, ability that is well reflected in the paradigm of the conditioning lesion. In fact, the robust regenerative response observed in juvenile neurons can be mediated, mainly, by the ability of these neurons to upregulate RAGs. In contrast, P4 and P10 neurons seem to be in a transition state, with a decline in the intrinsic growth capacity reminiscent of the embryonic stages but still not mature enough to activate a regenerative program in response to injury.

The incapacity of early postnatal neurons to activate the paradigmatic program than adult peripheral neurons upregulate can be inherent to the immaturity of neurons at these ages, but also to extrinsic factors, such the inflammatory response triggered by the injury, both at the distal stump and around neuronal somas. Despite it has been proposed that neonatal Schwann cells might fail to create a permissive environment for regeneration and participate in neuronal death (Low et al., 2003), in our previous work (Molina-Esteve et al., 2025), when characterizing Wallerian degeneration at different ages, we did not find evidence that that at early stages Schwann cells in the distal nerve stump failed to create a permissive environment for regeneration. In the other hand, macrophage infiltration at the DRG seems to contribute to the shift of injured neurons to a pro-regenerative state (Xin Lu and Richardson 1991; Xin Lu and Richardson 1993; Niemi et al., 2013; Kwon et al., 2013) and therefore, we cannot discard that differences in the inflammatory response between ages could had contribute to the final regenerative outcomes, both in vivo and in vitro. It is important to consider that, these conditioning lesion, despite being analysed in vitro, has been applied in vivo and, therefore, it partially dependent of this inflammatory response. In fact, neuronal Ccl2 by mediating neuron–macrophage interaction through monocytic Ccr2 receptor, has been proposed as a key element for conditioning sensory neurons (Niemi et al., 2013; Kwon et al., 2015). Ccl2 was significantly upregulated in DRG when injury was applied at P30 but not at early postnatal stages. Other chemokines that could compensate the lack of Ccl2 (Talsma et al., 2022) such as Ccl7 or Ccl2, also ligands of Ccr2, were neither upregulated at P4 and P10 after injury.

Ccr2, that was also markedly upregulated in P30 DRG after injury, is one of the main mediators of monocyte migration to tissues, and therefore, not typically expressed in neurons. But the fact that we analysed bulk RNA can lead to mixed responses from different populations of cells present in DRG, such as macrophages or satellite glial cells. In fact, when analysing specifically the translatome of motoneurons (Molina-Esteve et al., 2025), a larger activation of genes and pathways was observed, exposing the limitations of a non-specific tissue RNA analysis. But for technical reasons, mainly related with the low amount of specific RNA per subpopulation of sensory neurons, we chose a simpler design (whole DRG), that despite its constraints, allowed us to see a similar genetic profile between injured motoneurons and sensory neurons of the same age.

In conclusion, the age-dependent conditioning response observed in this study underscores the fundamental shift from an axonal growth program in neonatal neurons to a neurotransmitter state in mature ones (that retain the ability to shift to a pro-regenerative state if needed). The lack of upregulate RAGs in the injured neurons and the absence of conditioning lesion effect is in accordance with this appreciation.

## Funding

This work was funded by the projects PID2021-127626OB-I00 and PID2025-173721NB-I00 and from Ministerio de Asuntos económicos y Transformación Digital of Spain. The author’s research was also supported by funds from CIBERNED and TERCEL networks, co-funded by European Union (ERDF/ESF, “Investing in your future”).

## Acknowledgements

The authors appreciate the technical help of Neus Hernández. Mònica Espejo and Jessica Jaramillo.

## Declaration of Competing Interest

The authors declare no competing financial interests.

## Notes

### Competing Interest Statement

The authors have declared no competing interest.

