## Supplementary figures for "Outgrowth of Mouse Sensory Neurons is not Mediated by Regeneration Associated Genes in Early Postnatal Stages"

### Supplementary information

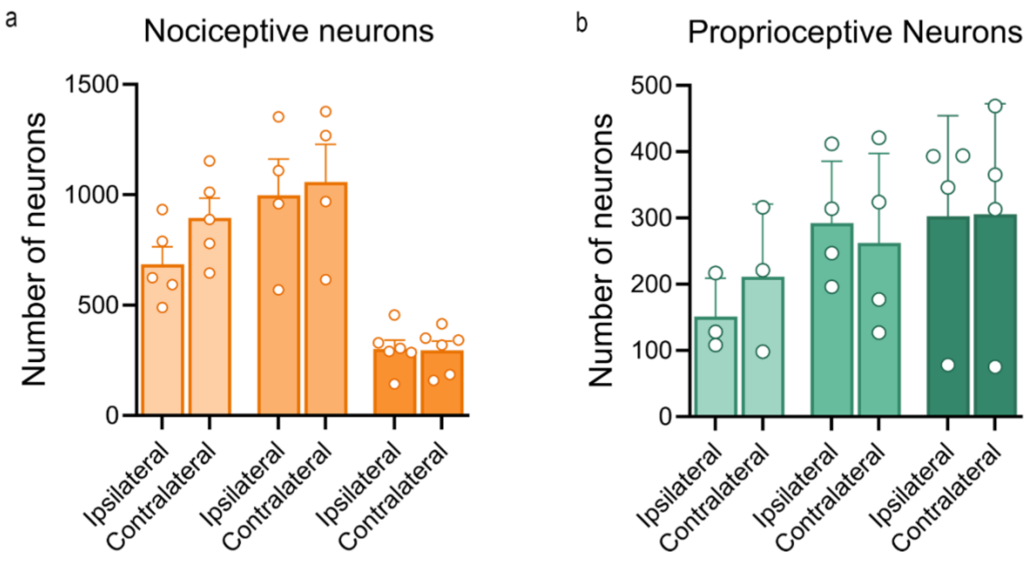


**Supplementary Figure 1**. (a) Quantification of the average number of PV$^{+}$ neurons in the L3-L5 DRGs on both the ipsilateral and contralateral sides at 14 dpi across different postnatal stages. Statistical analysis was performed via two-way ANOVA followed by Sidak's correction test. (b) Quantification of the average number of TrpU1$^{+}$ neurons in the L3-L5 DRGs on both the ipsilateral and contralateral sides at 14 dpi across different postnatal stages. Statistical analysis was performed via two-way ANOVA followed by Sidak's correction test.

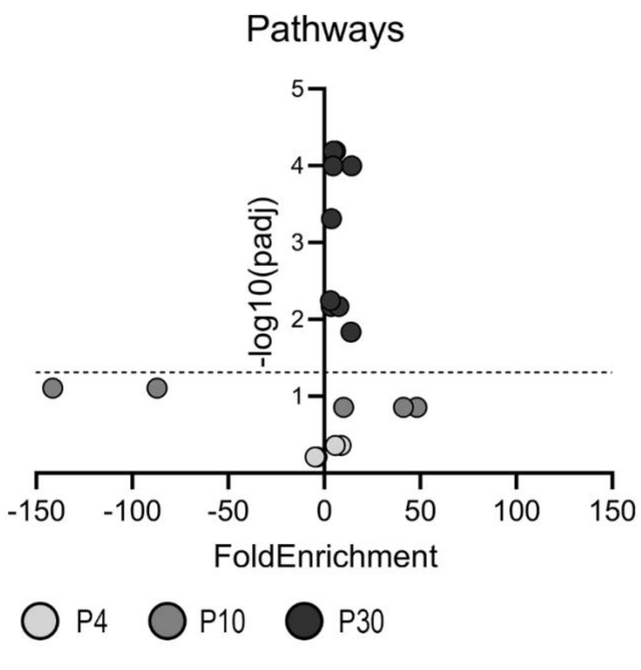


**Supplementary Figure 2**. Selected signalling pathways from animals injured at P4 (light grey circles), P10 (grey circles) and P30 (dark grey circles) showing fold enrichment and -log10(padj). Pathways below the dotted line correspond to those with *padj*>0.05 but *p-value*<0.05.

| P4 Crush vs P10 Control | | | | | |
|---|---|---|---|---|---|
| **Pathway** | **FoldEnrichment** | **pvalue** | **p.adjust** | **Gene Count** | **Key Genes** |
| Developmental Growth Involved in Morphogenesis | -3,98 | 0,000453 | 0,6165 | 9 | Nlgn3, Zeb2, Med12, Unc13a, Rtn4, Megf8, Nrp2, Csf1, Dclk1 |
| Neuron Projection Extension | -3,76 | 0,005410 | 0,6165 | 6 | Nlgn3, Unc13a, Rtn4, Megf8, Nrp2, Dclk1 |
| Axon Extension | -4,74 | 0,004160 | 0,6165 | 5 | Nlgn3, Rtn4, Megf8, Nrp2, Dclk1 |
| Cellular Response to Calcium Ion | 8,69 | 0,004980 | 0,4380 | 3 | Cpne1, Ryr3, Prkaa1 |
| Skeletal Muscle Organ Development | 5,75 | 0,001770 | 0,4380 | 5 | Prkaa1, Mylk2, Myoz1, Disp1, Acta1 |
| **P10 Crush vs P17 Control** | | | | | |
| **Pathway** | **FoldEnrichment** | **pvalue** | **p.adjust** | **Gene Count** | **Key Genes** |
| Regulation Of Inflammatory Response | 10,03 | 0,000546 | 0,1389 | 4 | Adam8, Pde2a, Sphk1, Ash1l |
| Negative Regulation of Neuron Projection Regeneration | -141,46 | 0,007050 | 0,0785 | 1 | Epha4 |
| Motor Neuron Axon Guidance | -87,05 | 0,011400 | 0,0785 | 1 | Epha4 |
| Cyclic Nucleotide Metabolic Process | 48,3 | 0,000766 | 0,1389 | 2 | Pde2a, Adcy4 |
| Regulation Of Acute Inflammatory Response | 41,26 | 0,001050 | 0,1389 | 2 | Adam8, Ash1l |
| **P30 Crush vs P30 Control** | | | | | |
| **Pathway** | **FoldEnrichment** | **pvalue** | **p.adjust** | **Gene Count** | **Key Genes** |
| Muscular cell development | 5,97 | 0,000000 | 0,0001 | 13 | Acta1, Mybpc2, Myom2, Myoz1, Tcap, Casq1 |
| Striated muscular cell development | 4,77 | 0,000000 | 0,0001 | 16 | Acta1, Ccl12, Mybpc2, Myom2, Myoz1, Tcap |
| Muscular cell differentiation | 3,74 | 0,000003 | 0,0005 | 17 | Atp2a1, Trdn, Casq1, Hrc |
| Calcium ion transport from ER | 14,32 | 0,000001 | 0,0001 | 7 | Atp2a1, Trdn, Casq1, Hrc |
| Calcium ion homeostasis | 3,63 | 0,000065 | 0,0068 | 13 | Atp2a1, Ccl12, Ccl7, Trdn, Casq1, Ccr5, Hrc, Ccr2, Cxcl9, Ccr1 |
| Humoral immune response | 4,66 | 0,000001 | 0,0002 | 15 | C1qa, C1qb, C1qc, Ccl12, Ccl7, Trf |
| Response to chemokines | 7,5 | 0,000041 | 0,0051 | 7 | Ccr5, Ccr2, Ccr1, Spi1 |
| Taxis | 3,17 | 0,000049 | 0,0057 | 16 | Ccl12, Ccl7, Ccr5, Ccr2, Cxcl9, Ccr1, Spi1 |
| Dendritic cell chemotaxis | 13,85 | 0,000182 | 0,0146 | 4 | Ccr5, Ccr2, Ccr1, Spi1 |

**Supplementary Table 1**. Significantly enriched biological processes in lumbar DRG (L3-L5) injured at P4, P10, and P30 and compared to age-matched uninjured controls. Pathways are ranked by statistical significance within each timepoint. Fold enrichment values indicate pathway activation (positive values) or suppression (negative values). Key genes represent the most relevant genes within each enriched pathway.

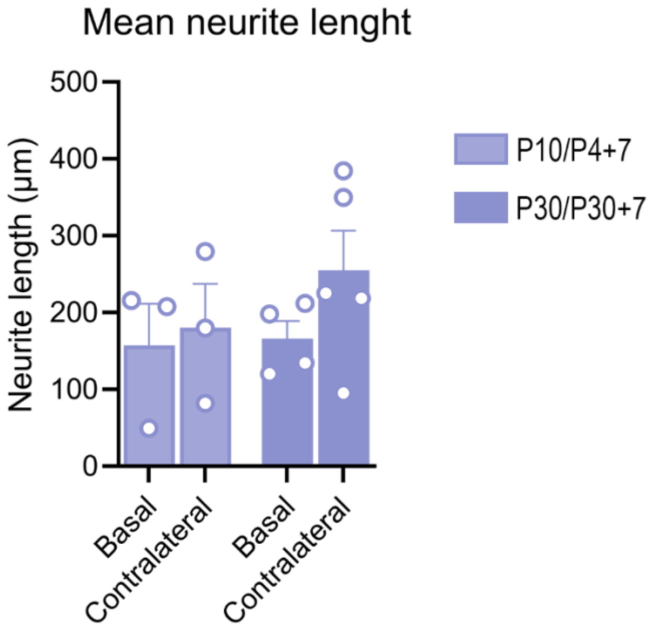


**Supplementary Figure 3**. Mean neurite length of neurons cultured in basal conditions (P10 and P30) neurons 7dpi from the contralateral uninjured side of mice injured at P4 and P30. Statistical analyses were performed using two-way ANOVA with Sidak's post hoc test.
